# Antiretroviral therapy in the peripartum period impairs post pregnancy cardiac reverse remodeling with early signs of heart failure with preserved ejection fraction

**DOI:** 10.1101/2025.09.11.675628

**Authors:** Nicole Taube, Mateus P. Mori, Cristina A. Nadalutti, Sherry Coulter, Madeline Jeshurin, Julia House, Don A. Delker, Michelle C. Cora, Helen Cunny, Suramya Waidyanatha, Georgia K. Roberts, Vicki Sutherland, Dawn M. Fallacara, Grzegorz L. Cielsieski, Janine H. Santos

**Author notes:** Global Toxicology & Safety Pharmacology J&J Innovative Medicine, Johnson & Johnson, 1000 U.S. Route 202 South, Raritan, NJ 08869. Corresponding Author: Janine Hertzog Santos, PhD.

## Abstract

Cardiovascular (CV) disease in pregnancy is the leading cause of maternal mortality worldwide, often associated with the cardiometabolic remodeling needed for fetal growth. In healthy pregnancies these changes reverse postpartum with no consequence, but improper reverse remodeling increases long-term CV risk. Pregnant patients with HIV are at further risk, but it remains unclear whether this is due to chronic low viremia or administration of antiretrovirals (ART). ART used in pregnancy includes nucleoside reverse transcriptase inhibitors (NRTIs), which have known mitochondrial toxicity, and may impact cardiac function. Here, we tested the hypothesis that NRTI-containing ART negatively impacts the maternal heart. Using rats, we show that pregnancy-associated reverse remodeling was impaired by ART. Molecularly, we found mitochondrial changes in the heart and in the liver, leading to dyslipidemia that collectively affected the heart. Our data suggest caution in the choice of ART prescribed for women with HIV during pregnancy and highlight the importance of studies focusing on maternal health.

## Introduction

Pregnancy is a dynamic process associated with significant cardiometabolic changes. During gestation, the increased hemodynamic load leads to cardiac hypertrophy that is accompanied by increased cardiac output and decreased systemic vascular resistance^1^. Metabolically, the mother switches carbon sources to include more lipids or ketone bodies favoring glucose uptake by the growing fetus^2^. Unlike in pathological hypertrophy, these physiological changes reverse after pregnancy – a process termed reverse remodeling (RR) that typically initiates a few weeks after birth, pending lactation, and is largely completed within 6 months post-delivery^3^. Maternal inability to adapt to these changes can expose underlying or previously silent conditions, including cardiac pathology that may persist postpartum. Owing to these, cardiovascular disease (CVD) in pregnancy is the leading cause of maternal mortality, including in North America^4^. Surprisingly, the mechanisms underlying these profound changes in the maternal cardiometabolic system, including the signals that drive RR, remain elusive. CVD in pregnancy is likely to be further compounded by environmental factors and comorbidities, such as chronic infections. In this context, CVD has also become a leading cause of morbidity and mortality in pregnant patients with HIV, who face additional physiological challenges due to the chronic viremia and the continued exposure to antiretrovirals (ART)^5^. The extent to which the presence of the virus and/or of ART further contribute to CVD in the pregnant HIV population is unknown.

Estimates indicate that 50% of the ∼40 million HIV^+^ population worldwide are women^6,7^. About 1.2 million women with HIV give birth globally per year, in the US alone the annual birth rate of women with HIV is estimated to be around 7,000^8^. HIV is currently a chronic condition with patients having relatively normal expected lifespan. Nevertheless, it remains imperative to understand the cumulative side effects of chronic ART exposure – particularly in the perinatal/peripartum period – given patients’ need to take these drugs throughout life. Notably, nucleotide reverse transcriptase inhibitors (NRTIs), which are known to negatively impact mitochondrial function by depleting mitochondrial DNA (mtDNA)^9^, remain a staple of HIV therapy, including during pregnancy where current treatment guidelines indicate the use of combination therapy with at least two NRTIs^10^. Given mitochondria are responsible for 90% of the ATP produced in cardiomyocytes and occupy 40% of their volume, one would predict adverse CV outcomes in pregnant patients taking NRTI-containing ART. Nevertheless, studies on pregnant women living with HIV or animal models have primarily focused on the effects of HIV/AIDS and its management in the offspring^9^, leaving an informational void about the off-target effects of the drugs on maternal health. By continuously exposing adult female rats from gestational day 6 (GD6) to postpartum day 120 (PPD120) to the NRTIs abacavir (ABC) and lamivudine (3TC) in combination with the integrase inhibitor dolutegravir (DTG), herein we characterized the effects of these drugs on cardiometabolic changes associated with pregnancy and RR. Our data revealed effects in the heart and liver that contribute to blunted RR in the postpartum period, collectively highlighting not only the need for further studies but the potential effects of NRTI-containing therapies in a vulnerable population.

## Results

### cART during the peripartum period does not affect pregnancy outcomes

A diagram of the experimental design is presented in Fig. 1. Briefly, dams were continuously exposed to 2 doses of abacavir/dolutegravir/lamivudine, 150/72.5/25mg/Kg or 300/1250/50mg/Kg, referred herein as low or high dose (LD or HD) combination ART (cART), from gestational day 6 (GD6) to postpartum day 120 (PPD120). These doses are equivalent to about 2.4X and 4.8X, respectively, the abacavir and lamivudine prescribed to a 60Kg patient while about 10 and 20X that of dolutegravir^11^. Cardiac assessments were done followed by tissue harvest at postpartum day 21 (PPD21) and PPD120 (Fig. 1A). These timepoints were chosen as the end of lactation (PPD21) and three months later (PPD120) to encompass the reverse remodeling period. No HIV virus or viral proteins were present under our experimental conditions, allowing us to dissect the effects of the drugs themselves. Pregnancy outcomes were generally similar between the groups, including in reproductive performance, fertility and fecundity, gestation length, litter size and numbers as well as clinical observations (Supplementary Data 1A). There were no significant changes in gross necropsy findings, including in the heart, kidneys, liver, lungs and mesenteric rolls (Supplementary Data 1B). No changes in food consumption or body weight were observed during pregnancy (Fig. 1B and C). While food intake was not changed in the postpartum period (Fig. 1D), cART-treated animals gained significantly more weight over time, which started at weaning (PPD21, Fig. 1E, F). Heart size also increased by treatment (Fig. 1G), but it was not different than controls if accounted for the increased body weight (Fig. 1H). Thus, cART treatment showed no signs of overt toxicity when administered during the time encompassing pregnancy, lactation and the 3 months that followed. This is consistent with previous studies, although clinical safety data on pregnant patients is sparse particularly when using the combined therapy^12^.

**Fig. 1:**
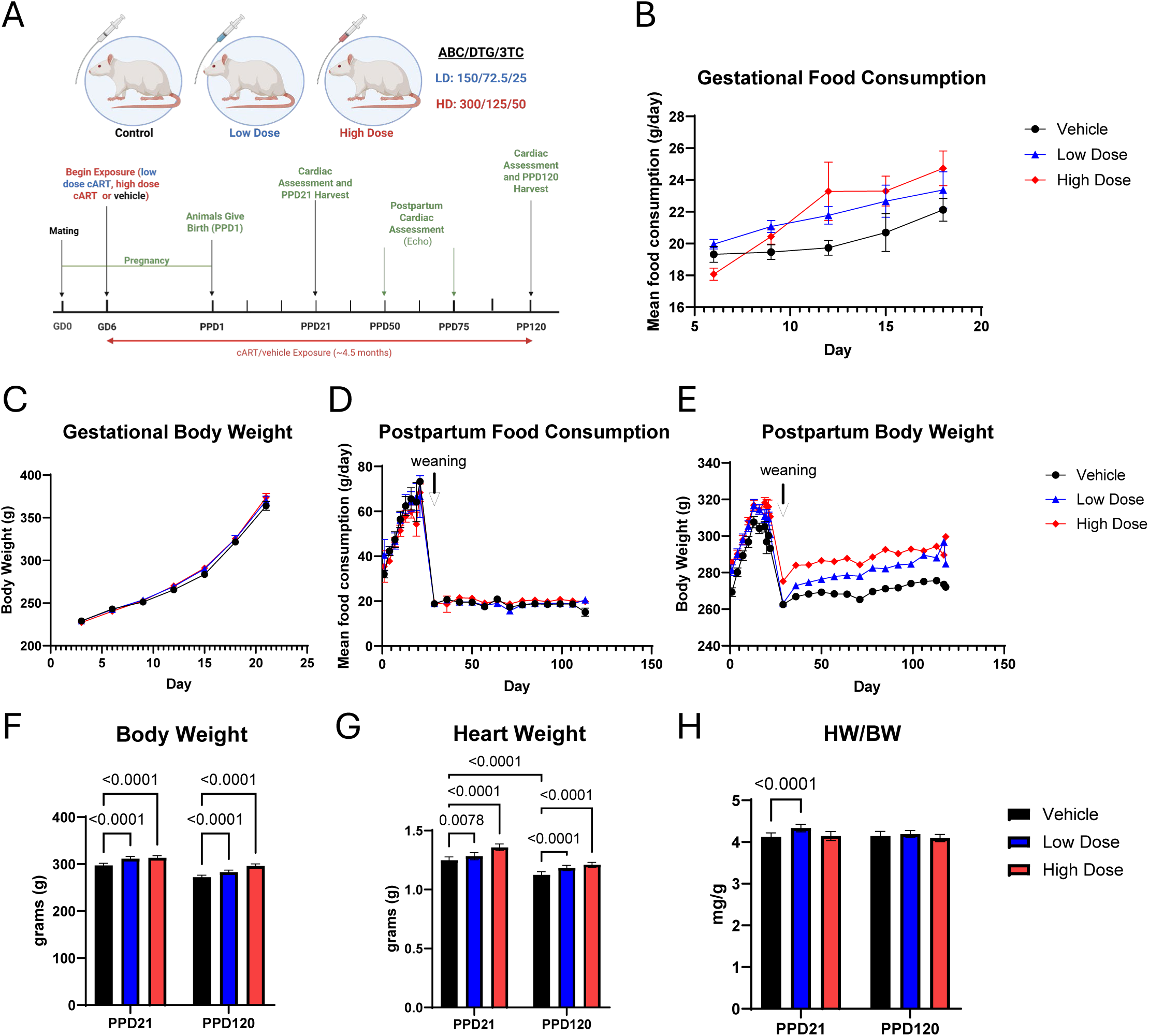
Peripartum ABC/DTG/3TC exposure increases body weight and heart weight in Sprague-Dawley female rats. (**A**) Diagram of animal exposure and timeline. Gestational food consumption (**B**) and body weight (**C**) over time (n=21-22 animals/group). Postpartum food consumption (**D**) and body weight (**E**) over time (n=21-22 animals/group). Body weight (**F**), heart weight (**G**) and heart weight normalized to body weight (**H**) at PPD21 and PPD120. Analyzed via two-way ANOVA, data shown as mean ± SEM. (n=11-12 animals/group/timepoint).

### Differential effects of cART on cardiac function during the peripartum period

We next characterized the effects of cART on CV function by echocardiography (ECHO). We used data from PPD21 to reflect pregnancy-induced changes given those initiate early in pregnancy and do not start reversing prior to weaning; data from PPD120 was used to inform on post-pregnancy physiology, including RR. Increased cardiac output (CO), stroke volume (SV), ejection fraction (EF) and fractional shortening (FS) are expected changes associated with pregnancy, which were significantly increased by cART at the HD at PPD21 relative to the controls (Fig. 2A-D). Heart rate (HR) was not altered under baseline measurements (Supplementary Data 2A), indicating that increased CO was driven by changes in SV. Similarly, animals on the HD showed changes in structural measures at PPD21 that were beyond the vehicle control, including increased left ventricular (LV) mass and posterior and anterior wall thickness as well as decreased systolic volume, which are consistent with physiological hypertrophy observed in pregnancy (Fig. 2E-I). Surprisingly, however, these parameters did not fully return to pre-partum physiology at PPD120, including in the LD group, with significant differences relative to the vehicle-controls still identified for CO. SV, EF, FS (Fig. 2A-D, F). These data indicate that the heart of the treated animals continued to have increased function at 4 months postpartum, despite no longer being pregnant, a period in which RR should have occurred. Notably, indication of RR was found in the control animals as early as 4 weeks post-weaning, but it was significantly blunted in the treated counterparts (Supplementary Data 2A). It is possible that the heart continued to work harder because of the increased size of the treated animals, which by the end of the experiments (PPD120) weighed essentially the same as the control animals at the end of lactation (PPD21) (Fig. 1E and F). Alternatively, but not mutually exclusive, they may reflect failure to initiate or complete RR.

**Fig. 2:**
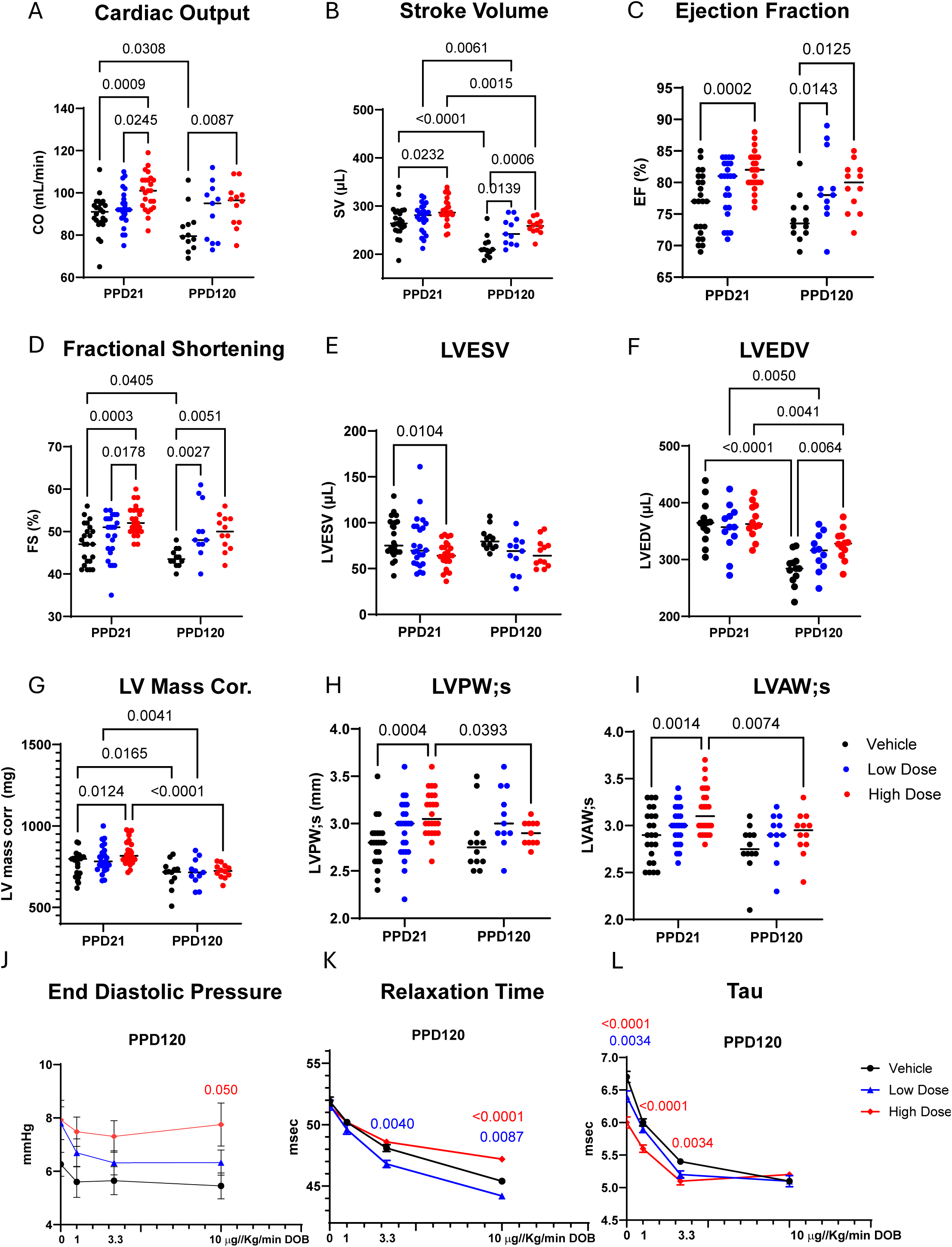
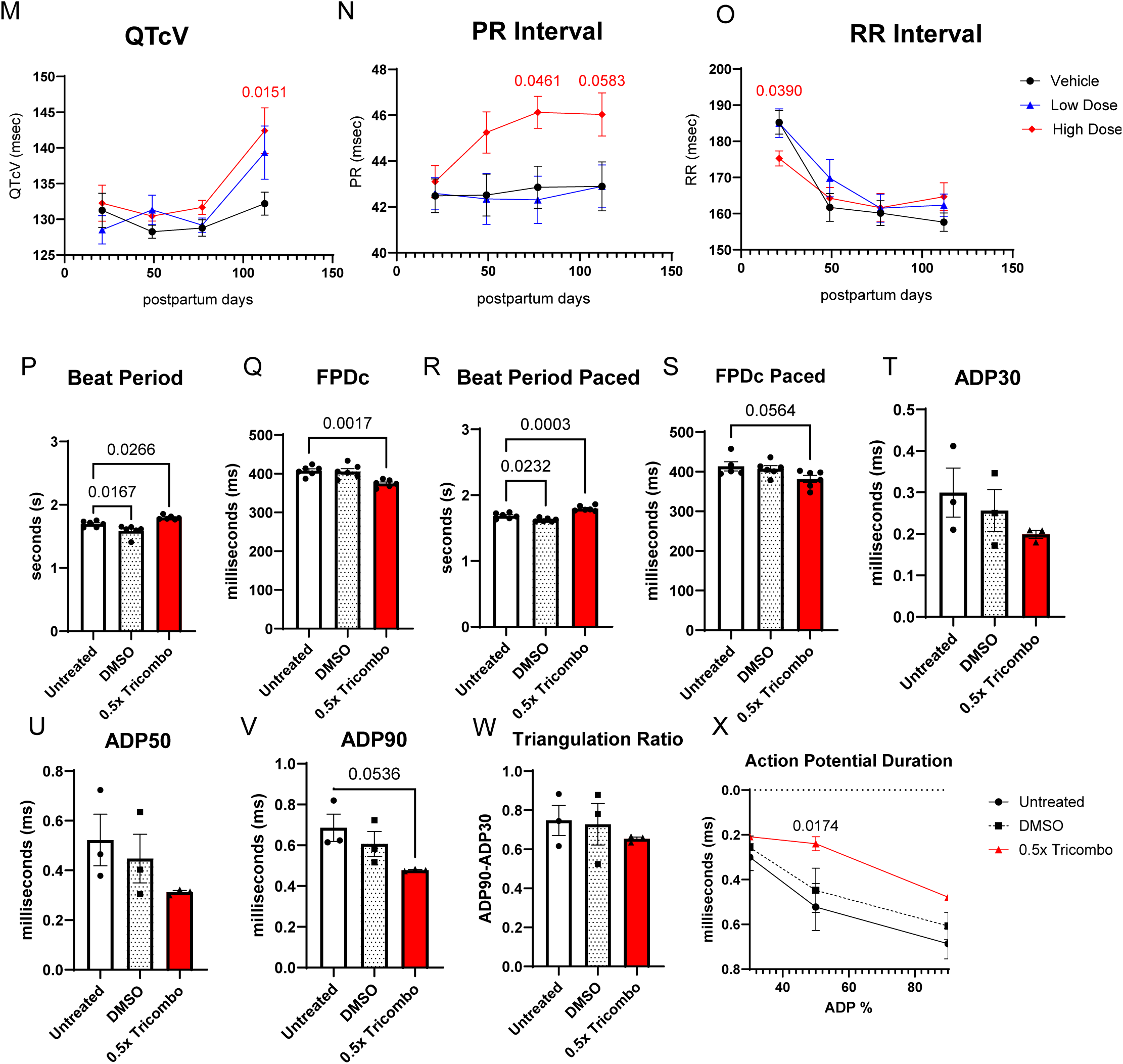
ABC/DTG/3TC administration alters contractile and electrophysiological function postpartum. Echocardiography measurements of vehicle, low and high dose ABC/DTG/3TC-treated dams at PPD21 (n=21-24/group) and PPD120 (n=12/group) for (**A**) cardiac output (CO), (**B**) stroke volume (SV), (**C**) ejection fraction (EF), (**D**) fractional shortening (FS), (**E**) left ventricular end systolic volume (LVESV), (**F**) left ventricular end diastolic volume (LVEDV), (**G**) left ventricular mass, (**H**) left ventricular posterior wall in systole (LVPW;s), (**I**) left ventricular anterior wall in systole (LVAW;s). Analyzed via two-way ANOVA with Tukey’s multiple comparisons (n=12-24 rats/group). (**J**) End diastolic pressure at PPD120, relaxation time (**K**), and tau (**L**) measured under dobutamine challenge. Analyzed via two-way ANOVA with Tukey’s multiple comparisons (n=12 rats/group), p-values shown as group vs. vehicle. Electrocardiogram measurements of vehicle, low and high dose ABC/DTG/3TC dams at PPD21 and PPD120 for (**M**) QTcV., (**N**), PR interval and (**O**) RR interval. Analyzed via two-way ANOVA with Tukey’s multiple comparisons (n=12 rats/group). Multielectrode array measurements for human iPSC cardiomyocytes dosed for 7 days with ABC/DTG/3TC including (**P**) beat period, (**Q**), field potential duration corrected for beat period, (**R**) paced beat period, (**S**) paced field potential duration corrected for beat period, (**T**) action potential duration 30%, (**U**) action potential duration 50%, (**V**) action potential duration 90%, and (**W**) triangulation ratio. (**X**) Action potential duration graphed over time. Analyzed via one-way ANOVA, data shown as mean ± SEM (n=6 wells/group).

We then submitted animals to a dobutamine challenge. Dobutamine is a β-adrenergic agonist that stimulates contractility and heart rate, informing on how the heart responds to sympathetic nervous system stimulation. Several significant differences were observed between the treated and vehicle-controls groups, with increased severity found in the HD-treated animals (Supplementary Data 2B). Notably, we found increased end diastolic pressure and increased relaxation time at PPD120 (Fig. 2J, K) – features often associated with diastolic dysfunction^13^ – in the HD-exposed animals. While initially we also observed a decrease in tau in this group compared to controls, as dobutamine concentration increased this difference was mitigated (Fig. 2L), indicating a reduced ability to respond to stress, or decreased contractile reserve, that is also seen in diastolic dysfunction^14^. Lastly, we found changes in baseline measurements of contractility, including decreased contraction time at both time points and increased contraction index (Supplementary Data 2B). The milder effects on the LD-treated animals indicate dose-dependency in the severity of the phenotypes. No significant difference in blood pressure was observed between the groups (Supplementary Data 2C).

We also performed electrocardiograms (ECGs) to define any effects on electrical signaling in the heart, which revealed increases in QT (at PPD120) and PR intervals, and a decrease in the RR interval at PPD21 that was significant in the HD-treated animals (Fig. 2M-O). No changes in QRS were observed (Supplementary Data 2D). An increase in QT indicates longer time between depolarization and repolarization of the left ventricle whereas that in PR reflects delayed electrical signals going from the sinoatrial node to the left ventricle. While these indicate that cART may increase the risk of arrhythmias or atrial fibrillation in the animals, measuring and interpreting ECG data from rodents is challenging because it can often be influenced by anesthetics and, given intrinsic differences to humans in ion channel expression, be of questionable translatability. To gain more insights into this, we treated iPSC-derived human cardiomyocytes in culture with the drugs, analyzing different ECG-relevant parameters using an impedance-based multielectrode array (MEA). Cells were exposed to doses equivalent to half (0.5X) of their concentration in the plasma, which is closer to the amounts in the heart (see below). Control experiments demonstrated that cells created a beating syncytium and were viable when kept in culture for up to 14 days. As beating patterns decreased and signs of drug toxicity increased by day 14, experiments herein were done in 7-day cardiomyocyte cultures.

About 50,000 cells per well were exposed to the drugs for 7 consecutive days, when field potential duration (FPD) and action potential duration (APD) were recorded in non-paced or paced cells. Pacing was used to synchronize the population so that any effects identified would be unrelated to spontaneous beating, and FPD was corrected to account for beat period, which is known to correlate to FPD. The FPD reflects the timing between depolarization and repolarization of all cells within each well, while the APD reports on the action potential of individual cells. We measured APD at 30%, 50% and 90% duration to examine how the APD changes over time, and to determine the triangulation ratio that informs on the repolarization rate of action potentials (ratio of APD50 to 90). We found that treated cells had increased beat period and decreased FPD whether the cells were or not paced (Figs. 2P-S). We also found reduced APD with cART exposure (Figs. 2T-X), together reproducing ECG changes observed in the rats, including those associated with QT prolongation. These collective findings indicate that cART exposure in the peripartum period leads to subclinical features of diastolic dysfunction, including altered electrophysiological properties and hypertrophy, that accompanies the blunted cardiac RR identified 4 months postpartum.

### cART alters cardiac mitochondrial DNA content and structure

The exact timing and molecular mechanisms driving pregnancy-associated RR remain poorly understood, but studies show that hormonal and metabolic rewiring, including of mitochondria, are involved^2, 15^. As two out of the three drugs used in our cART are NRTIs known to cause mitochondrial DNA (mtDNA) depletion^16^, we hypothesized that changes in mtDNA content and metabolism may be associated with the cardiac effects observed. To test this, we first determined the effects of the drugs on mtDNA content. We isolated heart genomic DNA and used primers covering different mtDNA regions (Fig. 3A). Data were normalized to a nuclear gene (*18S*) and, given similar trends (Supplementary Data 3A), were pooled to report on the entire genome as the average fold change of all primers for each animal. The HD group showed decreased mtDNA content at PPD21, with no changes observed between groups at PPD120 (Fig. 3B). Data from ND4L, which resides within the site of the common deletion^17^, was normalized to ND1 and used as a proxy for the occurrence of mtDNA deletions. Loss of the ND4L signal relative to ND1 was significant only at the HD group animals at PPD120 (Fig. 3C), suggesting the presence of mutated genomes missing that locus in those animals. An increased steady state of deletion-bearing mitochondrial genomes could trigger mtDNA biogenesis as a compensatory response, which can explain the lack of mtDNA content changes at PPD120 despite the continued presence of the drugs (Fig. 3B). Consistent with this, levels of 7S DNA, a tri-strand intermediate that originates during initiation of mtDNA replication that can be used as proxy for mtDNA biosynthesis, was found to be increased in the HD-treated hearts (Fig. 3D). When looking at mtDNA content as a function of time (pregnancy vs post-pregnancy physiology), we found lower mtDNA copy number between PPD21 and PPD120 in vehicle and LD animals, but not in the HD group which showed an increase in mtDNA abundance (Fig. 3E). Thus, the drugs had effects on mtDNA content that went beyond those associated with the physiological cardiac RR that occurs over time.

**Fig. 3:**
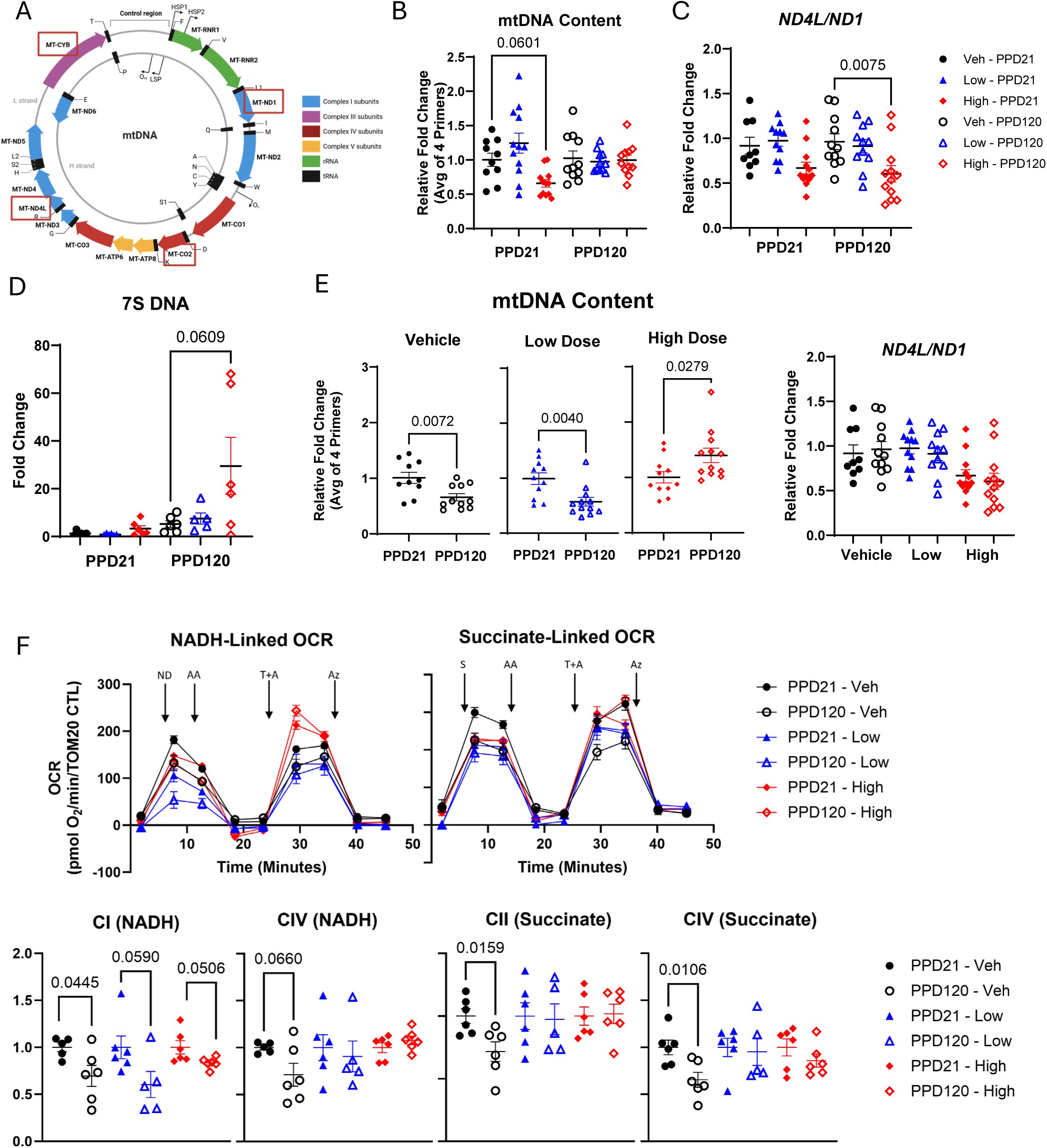
Mitochondrial DNA content and function are altered in the maternal heart by cART exposure during and after pregnancy. (**A**) Diagram of the mammalian mitochondrial genome with the location of genes analyzed indicated with purple rectangles. (**B**) mtDNA content as pooled data from individual primers for each animal relative to the nuclear genes coding for 18S, (**C**) *ND4L/ND1*, and (**D**) 7S DNA measured via qPCR at PPD21 and PPD120, analyzed within timepoints across dose groups. (**E**) mtDNA content and *ND4L/ND1* analyzed across timepoints within dose group. Analyzed via one-way ANOVA, data shown as mean ± SEM (n=12 rats/group). (**F**) Oxygen Consumption Rates (OCR) traces measured in lysates from frozen heart tissue in PPD21 and PPD120 within timepoint and (**G**) and PPD120 across timepoints using the Seahorse Flux Analyzer. OCR traces are shown above with their graphical representation depicted below. Results were analyzed via t-test, data shown as mean ± SEM (n=6 rats/group).

To determine whether the above changes impacted mitochondrial function in ways we could biochemically detect, we adapted a protocol for analysis of oxygen consumption using frozen heart lysates^18^. No significant changes in respiration were observed in the treated groups compared to the vehicle control at PPD21 or PPD120 (Supplementary Data 3B), much like the hiPSC-derived cardiomyocytes (Supplementary Data 3C). However, if accounting for differences in mtDNA content overtime, oxygen consumption was significantly decreased between PPD21 and PPD120 in the vehicle and HD group, while it was borderline in the LD animals when using complex I substrates; it was significantly decreased in the control animals when using the complex II substrate, succinate (Fig. 3F). Together with the changes in mtDNA content and the apparent increases in deletions on the genome of the HD group, we interpret these results to suggest that while the vehicle and the LD-exposed animals have decreased mitochondrial function over time as a component of normal cardiac RR, the HD-exposed animals compensate the presence of deleted genomes by increasing content. Nevertheless, those newly synthesized mtDNA molecules do not appropriately support bioenergetics.

The half-life of cardiac mitochondria is estimated to be ∼20 days^19^. Thus, while at PPD21 ∼75% of the initial population of mitochondria would have turned over, by PPD120 all that content would have been replaced. Given the continued presence of the drugs throughout the experiments, the next question we addressed was whether they affected mitochondrial steady-state levels. We used capillary electrophoresis and antibodies for the nuclear DNA-encoded mitochondrial matrix protein succinate dehydrogenase A (SDHA) and for the mitochondrial outer membrane protein TOMM20 (translocator of the outer mitochondrial membrane 20) but found no changes in their levels (Supplementary Data 3D). Therefore, we conclude that effects of the NRTIs are primarily seen at the mtDNA level consistent with their role in inhibiting mtDNA replication.

We also used transmission electron microscopy (TEM) to characterize impacts on cardiac ultrastructure. We found no differences in µultrastructure between the pregnant and post-pregnant state in the vehicle control animals (representative image on Supplementary Data 3D and summarized pathologist report). Conversely, mitochondria and other features of the cardiomyocyte structure were morphologically altered in both doses, although the number of altered mitochondria and degree of overall ultrastructural changes were more pronounced as the cART dose increased (representative images on Supplementary Data Fig. 3D). Increased glycogen granules were noted in the HD (see summarized pathologist report). Altered glycogen metabolism in the heart is a feature of metabolic stress and has been associated with cardiomyopathy since under normal circumstances glycogen breakdown provides glucose that can be used for bioenergetics, an important response in the heart^20^. Thus, the presence of ART altered mtDNA content, mitochondrial function and cardiac µultrastructure in ways that differed from those found upon physiological remodeling as observed in the control animals.

### Transcriptional profiles reveal effects on cardiac mitochondrial and lipid metabolism

The phenotypes so far identified suggest that cART biochemically affects the heart, even when provided in low doses. As we previously showed that progressive mtDNA depletion altered the epigenetic and transcriptional landscapes of cells in culture before any detectable signs of mitochondrial dysfunction^21, 22^, we next profiled the heart transcriptome. We first established the physiological changes occurring between PPD21 and PPD120 in the vehicle-controls, then determined how the drugs affected those. We found 1,446 genes that changed between PPD21 and PPD120 in the control animals (Fig. 4A). In the LD-treated animals we found 1,123 genes that were differentially expressed over time, while 3,234 genes were identified in the HD; common to all 3 groups were 393 genes (Fig. 4A). These common genes enriched for fatty acid metabolism and collagen (Fig. 4B). Genes that overlapped uniquely between the vehicle and LD animals (106) involved the immune system and ERK signaling (Fig. 4C), which is interesting because ERK has been associated with physiological and pathological cardiac hypertrophy^23–25^. Only 296 genes were common between the LD and HD (Fig. 4A), and while these included collagen fibril organization and fatty acid metabolism, we did not find significant similarities associated with mitochondria (Supplementary Table 1).

**Fig. 4:**
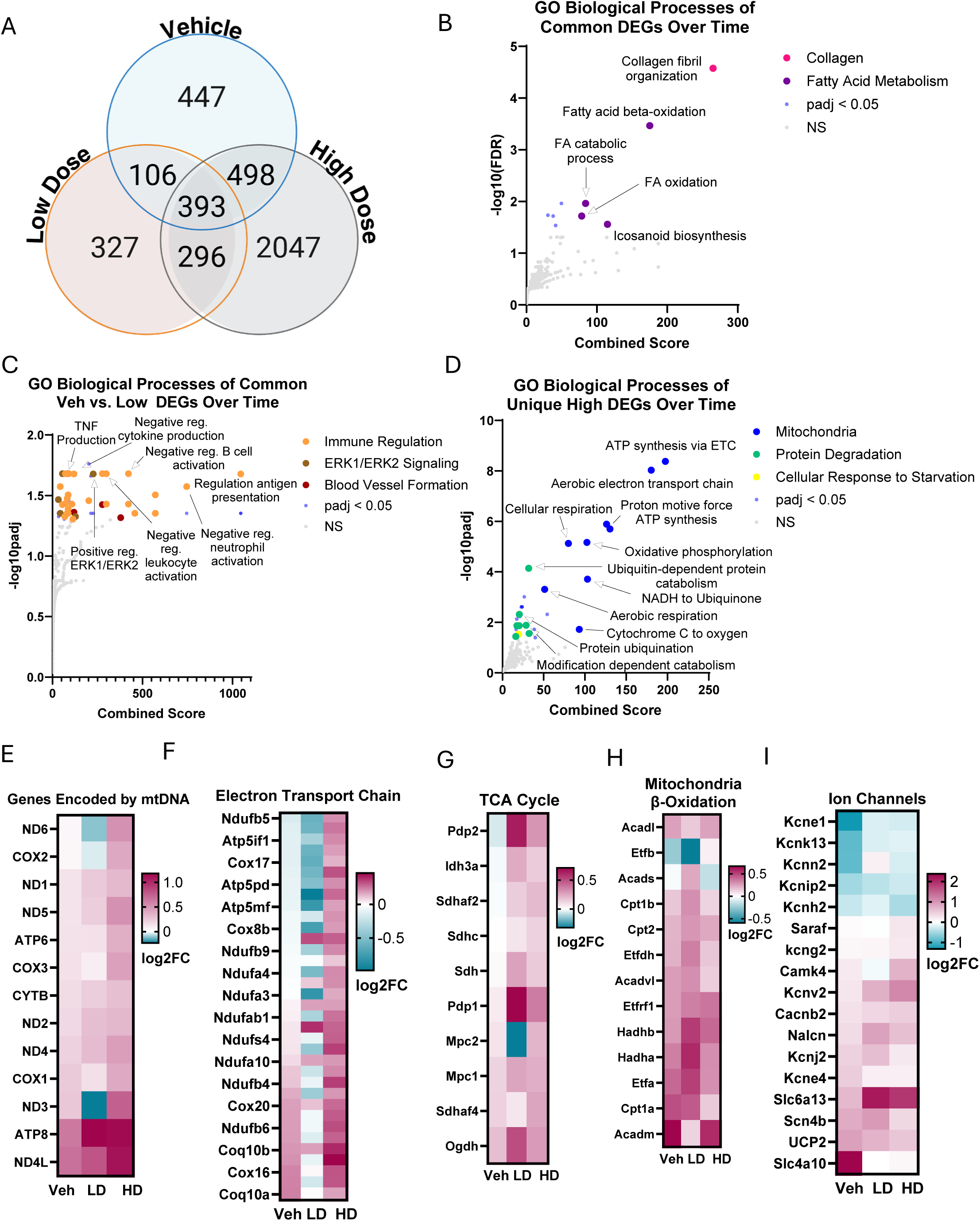
Significant transcriptional remodeling, including of mitochondria, follows cART exposure in cardiac tissue. (**A**) Venn diagram depicting the overlap of differentially expressed genes (DEGs) identified using RNA-seq in vehicle-, low dose- and high dose-treated animals. GO biological processes analyzed from DEGs (**B**) common to all three groups, (**C**) common to vehicle and low dose groups and (**D**) unique to high dose-treated animals over time (PPD21 vs. PPD120). Heatmap depicting fold differences in the expression of genes between (PPD21 vs. PPD120) associated with: (**E**) mitochondria, (**F**) electron transport chain, (**G**) TCA cycle, (**H**) mitochondria beta-oxidation and (I) ion channels (FDR < 0.05, n=6 rats/group/timepoint).

We next focused on the effects that were unique to each dose and found that the 327 genes that were solely changed in the LD-treated animals associated with calcium homeostasis (Supplementary Table 1) while the 2,047 uniquely altered in the HD enriched primarily for mitochondria and cellular response to starvation (Fig. 4D). The latter was noteworthy in view of the EM-observed glycogen accumulation in the heart, and suggest that despite ample fuel, the cardiomyocytes of the treated animals were not properly utilizing the available carbon sources. All transcripts from the mtDNA (Fig. 4E) and several nuclear-encoded ETC (electron transport chain) genes were upregulated in the HD-but not LD-treated animals (Fig. 4F) consistent with the changes observed at the mtDNA level (Fig. 3D). We also found differential expression of tricarboxylic acid (TCA) cycle and lipid oxidation genes based on the treatments, with mitochondrial beta oxidation seemingly activated in the LD but not in the HD when compared to the vehicle (Fig. 4G-H, Supplementary Table 1). Considering the differential effects on OXPHOS and beta oxidation with similarities in TCA cycle enzymes, these data suggest distinct fuel utilization by the mitochondria of cardiomyocytes based on the cART dose to which animals were exposed to. This is in line with the differential effects of the LD and HD on mitochondrial function over the same period (Fig. 3F). Interestingly, genes and pathways that we previously found to be causally linked to the broader effects of mtDNA depletion in cell culture^21, 22^ were also found to be differentially regulated in the treated animals (Supplementary Table 1), indicating important similarities in the broader transcriptional response to mtDNA depletion *in vitro* and *in vivo*.

Lastly, we found several genes coding for ion channels and solute carriers that were differentially regulated between the vehicle and the treated groups (Fig. 4I and Supplementary Table 1). Based on the directionality of changes of several of these ion channels, we would predict an increased in the intracellular Na^+^ pool, which could help explain the ECG phenotypes (Fig. 2O-Q). Notably, the largest differences in gene expression relative to vehicle-exposed controls involved transcripts from *KCNE1* (potassium voltage-gated channel subfamily E regulatory subunit 1) and the sodium bicarbonate exchanger *SLC4a10* (Fig. 4I) – both of which have been associated with long-QT syndrome^26^. Taken together, the RNA-seq data demonstrate the broad transcriptional remodeling that occurs in the heart as it transitions from pregnancy-associated to post-pregnant physiology. Importantly, it revealed that the presence of the drugs throughout this time had unique effects, including on mitochondria and their associated metabolism, that went well beyond those associated with physiological RR. Finally, the similarities and differences in the directionality and degree of the gene expression changes between the LD vs the HD collectively raise the intriguing possibility that the more severe cardiac changes observed in the latter are linked to the degree of cART effects on the mitochondria.

### Cardiac lipid metabolism is significantly modulated by cART

The significant transcriptional impact on mitochondria, prominent in the HD-treated animals, led us to profile the metabolome of the heart using mass spectrometry. The first observation was the detection of the drugs in cardiac tissue, which seemed to accumulate between PPD21 and PPD120 (Fig. 5A). As the same dose was delivered by gavage daily, we surmised this might reflect changes in blood volume between the pregnancy (PPD21) and the post-pregnancy state (at PPD120) ^27^. Little is known about the relationship between plasma and tissue concentration of ART, particularly when provided in the tri-combination prescribed to patients. Thus, we next took advantage of a method that we recently developed to detect abacavir, lamivudine and dolutegravir in biological samples ^28^ to gain insights into tissue levels. The liver was included in this analysis because it is the first site of metabolism and detoxification of exogenous drugs. Although we had heart from both doses and time points, we only had enough serum and liver available from the HD-treated animals at PPD120. Irrespective of this limitation, we gauged analysis of these samples was still of value to provide much needed comparative tissue level information. We found that the NRTIs abacavir and lamivudine were present in similar quantities in the liver, heart and serum (Fig. 5B, compare tan and violet bars), though they were less abundant in the heart if compared to the levels in the liver. Surprisingly, the integrase inhibitor dolutegravir (blue bar) was primarily detected in the serum with levels in the other tissues that were ∼ 10-fold less abundant (Fig. 5B), making it less likely to be a significant contributor to the cardiac phenotypes.

**Fig. 5:**
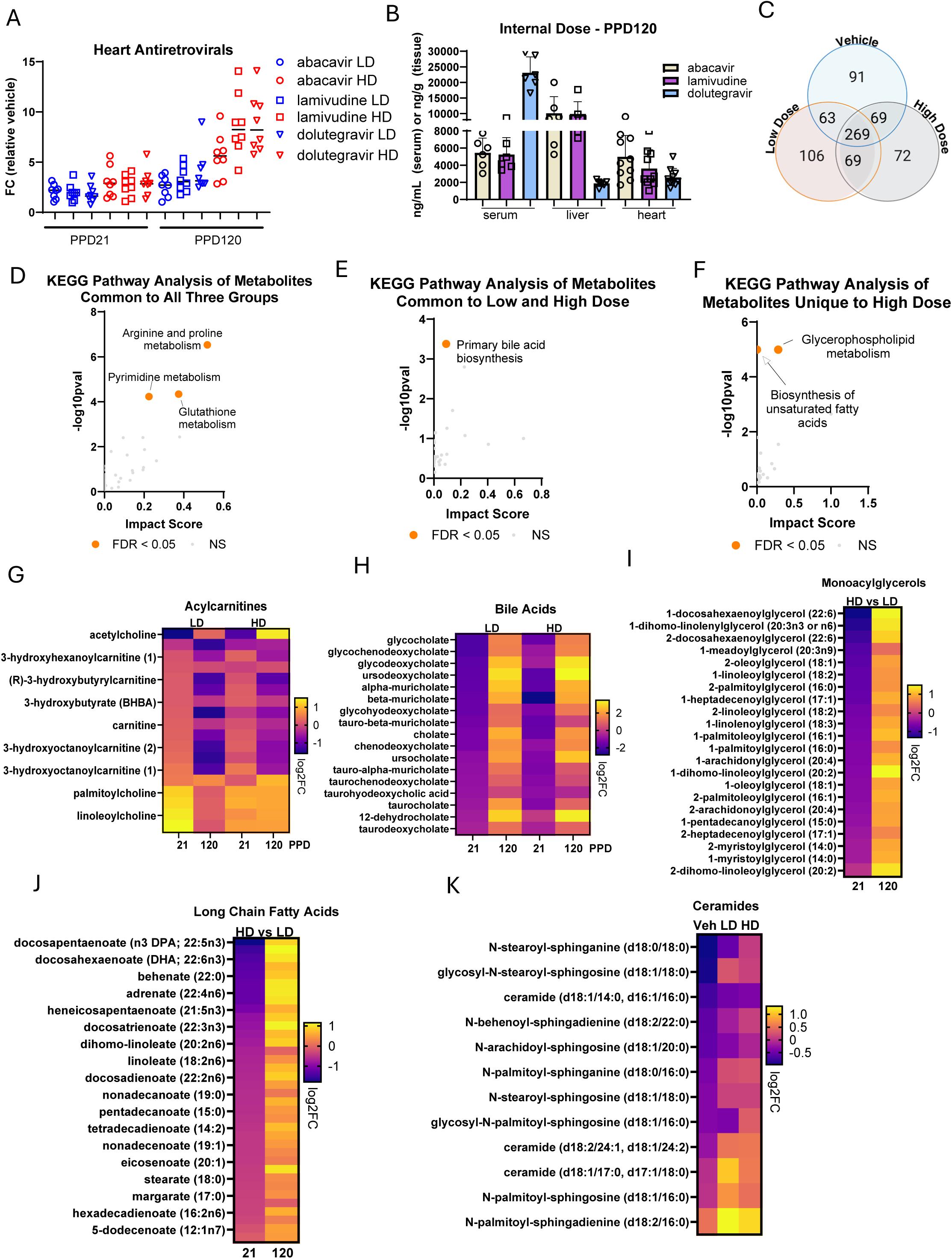
Metabolomics analysis indicates significant lipid remodeling in the heart, including the accumulation of bile acids, by cART. (**A**) Estimated levels of antiretrovirals detected in the heart by mass spectrometry at PPD21 and PPD120 in animals treated with the low and high dose cART. (**B**) Internal dose of antiretrovirals (ABC/DTG/3TC) in the serum, heart and liver of animals exposed to the high dose at PPD120. (**C**) Venn diagram depicting overlap of metabolites significantly changed over time (PPD21 vs. PPD120) in vehicle-, low dose- and high dose-treated animals. KEGG pathways analysis of metabolites (**D**) common to all three groups (**E**) common to low and high dose and (**F**) unique to high dose-exposed rats (FDR < 0.05, n=8 rats/group/timepoint). Heat maps depicting fold changes in low dose- and high dose-exposed animals at PPD21 and PPD120 for (**G**) acyl-carnitines and (**H**) bile acids. Heat map depicting difference in levels of metabolites (as fold change) when comparing high dose- vs low dose-treated animals at PPD21 and PPD120 for (**I**) monoacylglycerols and (**J**) long chain fatty acids. Heat map depicting the changes between PPD21 vs. PPD120 each individual group for (**K**) ceramides. (FDR < 0.05, n=8 rats/group/timepoint).

In the heart, specifically, we identified a total of 736 metabolites that were changed among the 3 groups, out of which 269 were common between them all (Fig. 5C). These shared metabolites enriched for amino acids, nucleic acids and redox metabolism (Fig. 5D, Supplementary Table 2), indicating that these are normal changes associated with physiological RR that were not affected by the drugs. Common to the two treated groups were only 69 metabolites that were involved, surprisingly, in primary bile acids biosynthesis (Fig. 5C and E) while unique to the HD were changes associated with additional types of lipids (Fig. 5C and F). Thus, unlike the transcriptional effects where the drugs had impact that went well beyond the normal changes associated with the transition from pregnancy (PPD21) to post-pregnancy physiology (PPD120), at the metabolome level the effect of the drugs was more muted. Nevertheless, some unique responses were noteworthy. For example, accumulation of acyl-carnitines and bile acids was a feature of both doses albeit the effects were more severe in the HD-treated animals at PPD120 (Fig. 5G and H). Similarly, changes in phospholipids were prominent in the treated animals (Supplementary Table 2) which is interesting given lysophospholipids are associated with inflammation in the context of HIV and have been linked to dyslipidemia and increased CVD in those patients ^29^. Conversely, monoacylglycerols (Fig. 5I) and long-chain fatty acids (Fig. 5J) decreased at PPD21 in the HD-relative to LD-treated animals but significantly increased in those animals by PPD120 (see also Supplementary Table 2). This accumulation of lipids can stem from increased lipid synthesis or decreased utilization, augmented lipolysis and/or altered gut lipid absorption. The increased levels of ceramides in the treated animals (Fig. 5K), a byproduct of the breakdown of triglycerides, supports the notion of increased lipolysis. Altered lipid metabolism, including through changes in lipolysis, has been shown to contribute to cardiac dysfunction^30^.

### cART-induced changes in liver metabolism impact systemic lipid levels

The detection of bile acids in the heart was unexpected because these lipids are generated in the liver and secondarily metabolized in the gut, after which they return to the liver via the portal vein to complete the enterohepatic circulation. However, a small amount of bile acids can escape this cycle, entering the systemic circulation thus reaching the heart^31^. As accumulation of bile acids in the heart has been shown to alter cardiac function, including through the inhibition of mitochondrial fatty acid beta oxidation^32^, these data lead to the intriguing possibility that, in addition to directly affecting the heart, the cART utilized in this study also impacted the liver and it is the combined effects in both organs that impacted cardiac function.

To test this, we next evaluated several liver-related clinical chemistry parameters. While we only had serum and liver available from animals at PPD120, we note that bile acids accumulated in the heart between PPD21 and PPD120 (Fig. 5H). Thus, we presumed that liver function, if altered, would be affected throughout the experimental timeline. Using serum from the animals we evaluated activity of the liver enzymes alkaline phosphatase (ALP), aspartate aminotransferase (ALT) and alanine aminotransferase (AST), but found no alterations to enzymes indicating no hepatocellular injury (Fig. 6A-C). These data demonstrated that the drugs did not cause overt liver damage. Conversely, we found mildly increased total bilirubin in the HD-treated group (Fig. 6D), although no significant changes in its conjugated form (direct bilirubin) (Fig. 6E), borderline significant increases in total circulating bile acids (Fig. 6F). These mild changes may indicate some degree of cholestasis (slowing of bile flow through the liver) and/or perturbation of normal bile acid metabolism. We additionally found increased total and high-density lipoprotein (HDL) cholesterol (Fig. 6G and H). No changes in LDL (Fig. 6I) or triglycerides (Fig. 6J) were identified. Notably, humans and rodents differ in cholesterol transport. While humans primarily transport cholesterol in low density lipoproteins (LDL), rodents transport primarily through HDL^33^. The primary reason is that rodents lack the enzyme cholesterol ester transferase protein (CETP)^34^, which facilitates the transfer of cholesterol and triglycerides between different lipoproteins^35^. Collectively, the lipid profiles show that changes in the local heart metabolome were paralleled by systemic alterations identified in the circulation.

**Fig. 6:**
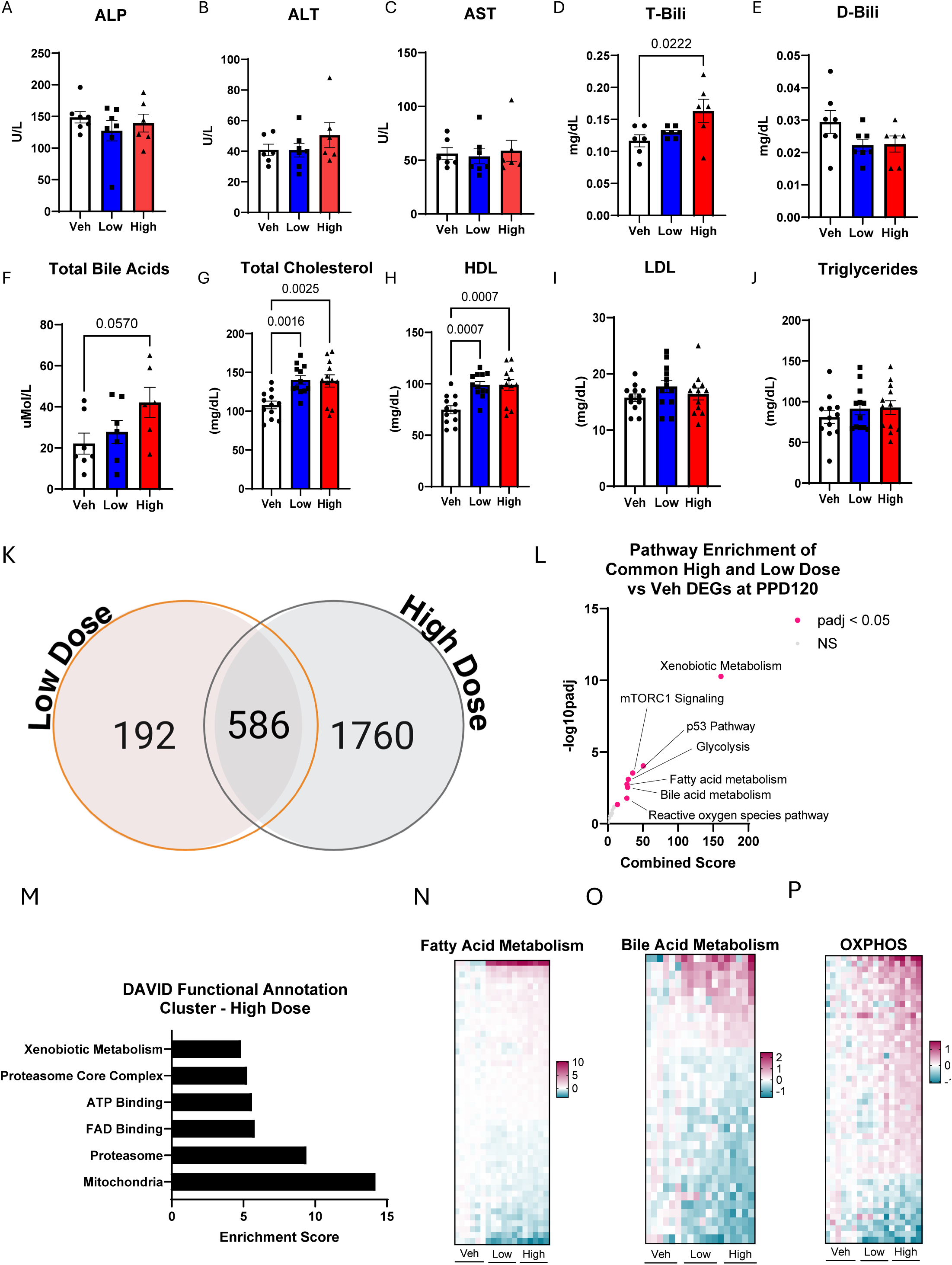
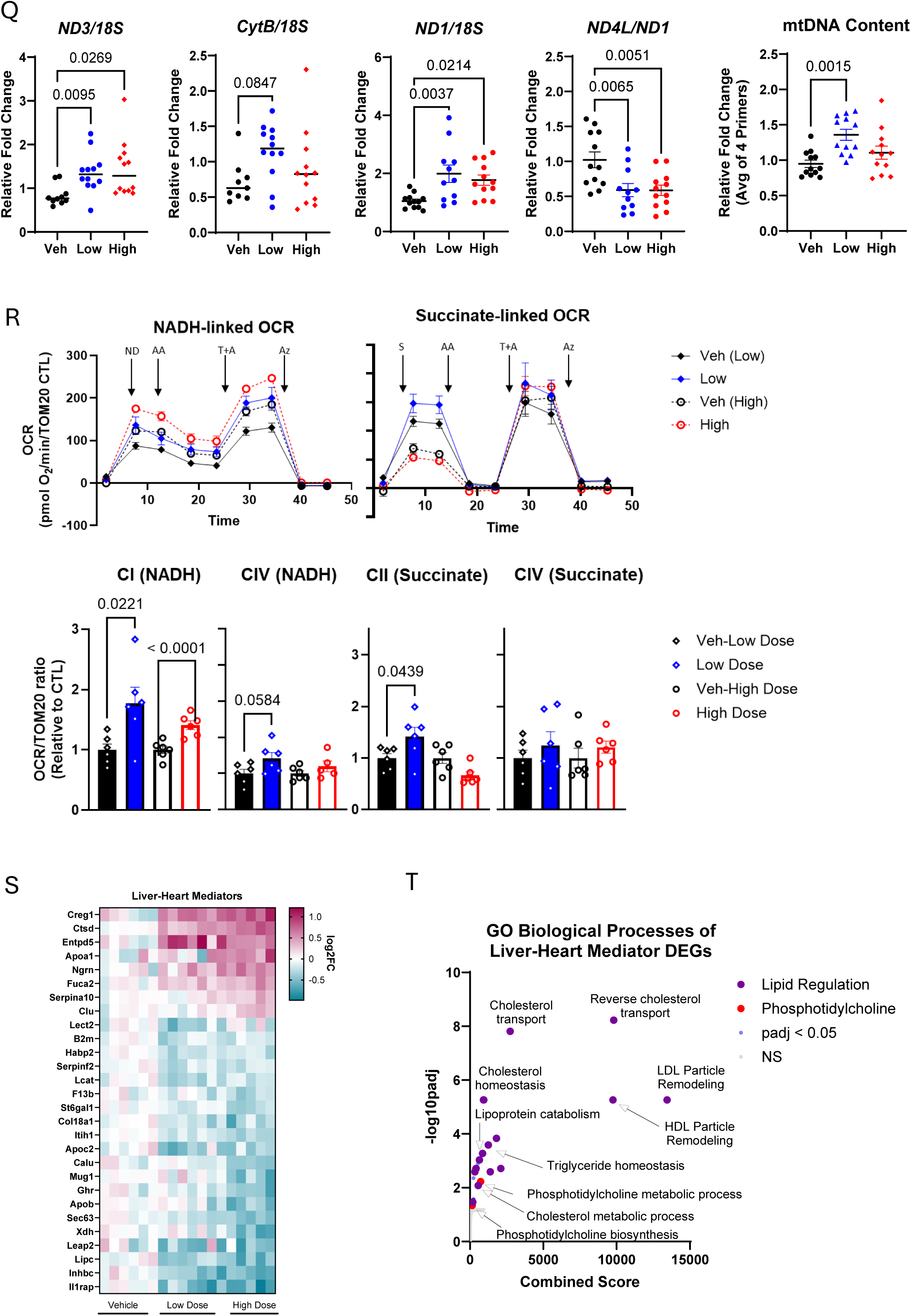
cART administration alters metabolism and mitochondria in the liver, leading to altered plasma lipid levels. Clinical chemistry measurements on serum collected at PPD120 for the HD group (**A**) ALP, (**B**) ALT, (**C**) AST, (**D**) total bilirubin, (**E**) direct bilirubin, (**F**) total bile acids, (**G**) total cholesterol, (**H**) HDL, (**I**), triglycerides and (**J**) LDL. Analyzed via one-way ANOVA with multiple comparisons (n=6-7 rats/group). (**K**) Venn Diagram of unique and overlapping liver genes differentially regulated by cART at the LD or HD relative to vehicle controls. (**L**) Pathway analysis of genes that were common between the two doses using MSigDB Hallmarks. (**M)** Functional annotation clustering (FAC) using DAVID based on RNA-seq from liver at PPD120 and heat maps showing differences as fold-changes relative to vehicle for (**N**) fatty acid metabolism, (**O**) bile acid metabolism and (**P**) oxidative phosphorylation (FDR < 0.05, n=6 rats/group). (**Q**) mtDNA content assessed by qPCR as in Fig. 2A. Analyzed via one-way ANOVA (n=12 livers/group). (**R**) OCR measured in frozen liver tissue from HD-treated animals at PPD120. Analyzed via t-test, data shown as mean ± SEM (n=6 livers/group). (**S**) Heatmap of liver-heart mediators identified via RNA-seq in liver as per Cao et al. (2022) ^36^. (**T**) GO biological processes pathway analysis using the identified liver-heart mediator genes shown in Q (FDR < 0.05, n=6 livers/group).

Next, we analyzed the transcriptome of the PPD120 livers using RNA-seq. Comparing LD and HD-treated livers to vehicle controls, we identified a total of 2,538 genes that were differentially expressed in the treated animals (Supplementary Table 3); about 586 were common to both groups while 192 were unique to the LD group and 1,760 to the HD (Fig. 6K). Pathway enrichment analysis using genes that were common to both LD-and HD-treated groups showed enrichment for xenobiotic metabolism, fatty acid and bile acid metabolism, in addition to glycolysis (Fig. 6L). Conversely, mitochondria represented the largest enrichment using the 1,760 genes unique to the HD (Fig. 6M) while no significant enriched pathways were found when using the 192 genes uniquely modulated by the LD (Supplementary Table 3). The change in transcription of fatty acids (Fig. 6N) and bile acid metabolism genes (Fig. 6O) in the liver tissue is consistent with the elevation in lipids and bile acids identified in the circulation. While the biggest fold differences were observed for fatty acid metabolism genes (Fig. 6N), 7% of genes in the dataset were associated with the mitochondria (Supplementary Table 3). Notably, 51 of these were specifically involved with oxidative phosphorylation (OXPHOS), suggesting alteration hepatic mitochondrial function (Fig. 6P). We found that mtDNA content was increased at PPD120 in the liver of treated animals while identifying decreased normalized ND4L amounts (Fig. 6Q). These data indicate that, like in the heart, the drugs activated mitochondrial biogenesis to compensate for damaged/mutated mtDNA. The effects on mtDNA, including on OXPHOS gene transcription, were paralleled by functional changes as judged by oxygen consumption from frozen mitochondrial-enriched lysates; complex II-driven respiration in the HD-treated livers - while tending to be lower than vehicle controls - did not reach statistical significance (Fig. 6R).

A recent study leveraged a bioinformatics approach that uses natural variation to find correlations that mediate organismal tissue crosstalk. Using the global transcriptome of the heart and the liver of 100 different diverse inbred strains of mice, they identified a set of 100 candidate liver genes to mediate the communication with the heart. By using several genetic and molecular approaches, they went on to show a causative association between excretion of a liver protein with heart failure with diastolic dysfunction and preserved ejection fraction ^36^. To get insights onto whether the changes in the liver transcriptome might influence cardiac function under our experimental conditions, we mined our liver RNA-seq data relative to the candidate gene list previously described ^36^. Remarkably, we found that 30 out of the 100 candidate genes were differentially regulated in the liver of the treated animals (Fig. 6S). Interestingly, the significantly enriched pathways associated with those genes primarily involved lipids (Fig. 6T). Taken together, these results indicate that cART-induced impairment of cardiac reverse remodeling may not be solely due to direct drug effects on the heart. Instead, our data suggest that perturbations in liver homeostasis, marked by altered lipid and bile acid metabolism, mitochondrial stress responses, and activation of xenobiotic and OXPHOS pathways, may contribute to a systemic environment that indirectly influences cardiac physiology – at least in the context of pregnancy-associated RR. While this supports the emerging concept of a liver-heart axis in drug-induced cardiometabolic dysfunction, our results take it a step further by implying that hepatic adaptation to mitochondrial stress could act as a mediator of long-term cardiovascular effects under chronic cART exposure.

## Discussion

In this study, we showed that maternal cardiac function, particularly postpartum RR, was impaired by the presence of abacavir/dolutegravir/lamivudine an antiretroviral combination that until June 2025 was the guideline medication for pregnant women living with HIV. We found that these effects correlated with altered mtDNA content and function and were associated with significant changes in lipid metabolism both in the heart and in the circulation - pointing to additional effects in the liver. While analysis of serum clinical chemistry did not identify any overt signs of liver injury, the transcriptomic data demonstrated significant mitochondrial and lipid remodeling. It also revealed the differential modulation of mediators previously described to connect liver function with diastolic dysfunction with preserved ejection fraction^36^ – phenotypes, albeit subclinical, observed in our animals.

Dyslipidemia is a common diagnosis in patients with HIV and has been linked to increased risk of CVD. While earlier epidemiological data suggested it was mostly due to the virus given that ART improved lipid profiles^37,38^, more recent studies have associated the exposure to ART with increased dyslipidemia; notably, lamivudine and abacavir were considered of low risk^37,38^. Mechanistically, how ART can lead to dyslipidemia is poorly understood but our transcriptional and metabolic data in the heart and the liver - in addition to internal dose measurements - strongly indicate that they are driven by the NRTIs and their effects on mitochondria. We note that mitochondrial complex activity decreased in the heart yet increased in the liver despite similar effects on mtDNA content. This divergence may reflect the differences in tissue-specific metabolic roles. Both heart and liver rely on fatty acid oxidation for ATP production. The liver, however, plays a central role in systemic metabolic regulation, producing glucose during fasting via gluconeogenesis, and fatty acids during feeding via *de novo* lipogenesis. Thus, changes in complex activity in the liver may reflect anabolic reprogramming or mitochondrial adaptations to altered lipid and bile acid metabolism induced by cART rather than increased oxidative demand.

The accumulation of acyl-carnitines and glycogen in the heart followed by the transcriptional response to cellular starvation suggest that the tissue is under metabolic stress, with neither fatty acid nor glucose being properly utilized. Further, the accumulation of free fatty acids, long chain fatty acids and monoacylglycerols in the heart, indicative of increased lipolysis, point to a mismatch between fuel availability and utilization. Interestingly, bile acids – found here to accumulate in the circulation and the heart - can also inhibit mitochondrial beta oxidation, providing yet another means through which mitochondrial activity can be negatively modulated under our experimental conditions. While it is unclear whether the identified bile acids impact mitochondrial cardiac function, our results on exposure of non-pregnant animals to the same cART regimen showed similar changes in circulating cholesterol and bilirubin but not in bile acids. Notably, those animals did not present with cardiac dysfunction other than minor hypertrophy (Marchese, Taube et al., in preparation). It remains to be defined if this indicates differential effects of cART in the liver or increased susceptibility provided by pregnancy itself, including through effects in the placenta, areas that currently under investigation in our laboratory.

Beyond these findings, several questions arise from our results. For example, the degree to which the effects of the drugs on the mitochondria were causative to the cardiac phenotypes and how these may mechanistically mediate the local (heart) and systemic (likely driven, in part, by the liver) lipid remodeling needs further investigation. The dose dependency of the effects on mitochondria and on the cardiac phenotypes suggest that a causative relationship between those two is plausible. It is also unclear whether the post-partum cardiac changes reflected either impaired or delayed pregnancy-associated RR given that our animals were only monitored for 4 months postpartum. It is possible that longer follow ups would demonstrate complete recovery or the maintenance of CV defects so long as the animals remain under cART exposure. Further studies are required to address these questions.

Irrespective, it is noteworthy that epidemiological studies have shown that increased HFpEF is a feature of women living with HIV, particularly those under NRTI-containing regimens^39^. It is also known that women who are living with HIV are at an increased risk of coronary artery disease with heightened activation of the immune response, despite ART viral suppression, and increased plaque deposits^40^. Although it is not clear whether these derive from off-target effects of the drugs or the presence of larger viral reservoirs in difficult to access tissues^40^, the absence of the virus or viral proteins under our experimental conditions suggest some contribution of the medications. Lastly, it was recently shown that women with ≥ 5 live births are at increased risk of developing future HFpEF, thus highlighting how cycles of physiological hypertrophy and RR concomitant with a second hit, such as cART exposure, may further increased the odds of women with HIV developing HF^41^.

In summary, our results point to a need for more work focusing on maternal, in addition to offspring health, in the population exposed to cART. While one significant limitation of our study was the lack of parallel analysis from human patients exposed to these drugs during pregnancy, we are currently exploring datasets that may help shed light into the translatability of our findings. The observed cardiac phenotypes in the animals were mild and subclinical, but they may represent a significant problem when patients are faced with additional stresses or environmental exposures in their lives, including those associated with comorbidities, medications and lifestyle choices.

## Materials and Methods

### Animals and cART treatment

Animal care and use were in accordance with the Public Health Service Policy on Humane Care and Use of Animals. All animal studies were conducted in an animal facility accredited by AAALAC International. Studies were approved by the Battelle (Columbus, OH) Animal Care and Use Committee (ACUC protocol T06055) and conducted in accordance with all relevant National Institutes of Health animal care and use policies and applicable federal, state, and local regulations and guidelines. Time-mated (F0) adult female Sprague Dawley rats, aged 11-14 weeks, were received from Envigo (Indianapolis, IN), housed individually during gestation and with their respective litters during lactation, until weaning (lactation day 21), after which they were group-housed. Animals were maintained under specific pathogen-free conditions in a 12h light cycle with water and food ad libitum. Time-mated rats were dosed daily via oral gavage (5 mL/Kg) starting at gestational day 6 (GD6) until about 13 weeks after weaning, totaling 4.5 months. Female rats were dosed with abacavir (ABC), dolutegravir (DTG) and lamivudine (3TC) in a combination ratio of 12:1:6, respectively, representative of clinical formulation. Formylations were prepared by RTI International (Durham, North Carolina). Females were dosed with either aqueous 0.2% methylcellulose/0.1% Tween 80 (vehicle), 150 ABC/12.75 DTG/75 3TC mg/Kg (low dose) or 300 ABC/25 DTG/150 3TC mg/Kg (high dose).

Cardiovascular assessments, electrocardiography (ECG) and echocardiography (ECHO), were performed in the F0 animals prior to weaning (∼PPD20), at 4 weeks (±7 days), at 8 weeks (±9 days), and at 13 weeks after weaning. Assessments within a target date were staggered based on assay throughput and selected to ensure adequate representation across dose groups. Once the animal order was established at the preweaning assessment, this order was repeated at the Week 4, Week 8, and Week 13 timepoints to ensure time between assessments for an individual animal was similar across all animals. Hemodynamic assessments were conducted as a terminal procedure, after the cardiovascular assessments were completed, for a subset of the F0 animals (n=12/group) at PPD20 and for remaining (n=12 per group) F0 animals at 13 weeks after weaning. Hemodynamic assessment was conducted by recording left ventricular and systemic pressures following a rising dose dobutamine challenge.

### Electrocardiogram

All animals were lightly anesthetized with isoflurane to effect (∼3%) via induction chamber and then maintained with ∼1.5 to 3% isoflurane with a cone mask for the conduct of cardiovascular data collection. Both ECG and ECHO were collected in the same session. Three electrodes were attached to animals for lead II ECG monitoring and recorded on a computer using the IOX Data Acquisition system (EMKA Technologies) The following ECG parameters were recorded and assessed: PQ Interval (msec), QRS Interval (msec), QT Interval (msec) and QTcV (QT corrected for heart rate using Van de Water’s formula) (msec).

### Transthoracic echocardiography

Following stabilization under anesthesia, left ventricular (LV) function and dimensions were monitored and recorded using a Vevo 3100 Ultrasound machine with a 25 to 55MHz probe (FUJIFILM VisualSonics Corporation, Canada). Transthoracic two-dimensional (2D) B-mode, as well as 2D-guided M-mode, echocardiography was obtained from the long- and short-axis views at the level of papillary muscles. A simultaneous single-lead (e.g., lead II) ECG was obtained and recorded for the determination of cardiac cycle intervals during ECHO evaluation with the ECHO images and cine loops. The recorded images and cine loops were exported and analyzed offline. The following ECHO parameters were recorded and assessed: heart rate (beats per minute [bpm]), systolic diameter (mm), diastolic diameter (mm), systolic volume (µL), diastolic volume (µL), stroke volume (µL), ejection fraction (%), fractional shortening (%), cardiac output (mL/min), left ventricular (LV) Mass (mg), LV Mass Corrected (mg), systolic LV anterior wall thickness (mm), diastolic LV anterior wall thickness (mm), systolic LV posterior wall thickness (mm) and diastolic LV posterior wall thickness (mm).

### Hemodynamic Assessment with Dobutamine Challenge

For the hemodynamic assessments, anesthesia was induced by isoflurane (up to 5%) via a gas chamber and maintained via face mask with 95% O_2_/5% CO_2_. Following anesthesia induction, the animal was shaved for vascular access and positioned in dorsal recumbency on a heated surgery table. Infiltration anesthesia at the incision site(s) (i.e., femoral triangle, jugular furrow, or ventral cervical region) was done with 2% lidocaine (∼0.05 mL diluted with 0.9% NaCl) administered prior to the instrumentations. A tracheostomy was performed for ventilatory support, and the animal was ventilated (∼100 breaths/min, ∼2.5 mL tidal volume with 95% O_2_/5% CO_2_) with an adjustable small animal ventilator (Harvard Apparatus). A suitable vein (left or right jugular or femoral vein) was isolated and cannulated for administration of dobutamine.

Transthoracic needle electrodes forming a single-lead ECG (e.g., lead II) were placed. To record arterial pressures, a 1.4F or 2F high-fidelity micromanometer catheter (Millar Instruments) was placed via a cut down and inserted into a femoral artery or carotid artery and advanced towards the aorta. To record LV pressures, the right carotid artery was isolated, dissected free from the surrounding tissue, and cannulated with a 1.4F or 2F high-fidelity micromanometer catheter (Millar Instruments). This catheter was advanced retrograde across the aortic valve and into the LV chamber. A vein (i.e., the other jugular or femoral vein) was isolated and cannulate with an appropriately sized IV catheter or polyethylene (PE) tubing for drug administrations, as necessary. Heparin (∼100 IU, IV) may be administered after the surgery. Following a baseline/hemodynamic stabilization period (up to 20 minutes), the overall cardiovascular responses were monitored for up to 120 minutes following baseline data collection.

Dobutamine challenges were administered by IV infusion for F_0_ dams by intraperitoneal (IP) injection for F_1_ pups in the Interim Cohort. Dobutamine was delivered in a dose escalation pattern at 0, 1, 3.33, and 10 µg/kg by IV infusion targeting 5-7 minutes at each dose. Hemodynamic recovery was allowed (up to 10 minutes) between each challenge. All animals were euthanized at the end of the experiment while they were under a surgical anesthetic plane. The following HEMO parameters were recorded and assessed, for the left ventricle: Left Ventricle End Diastolic Pressure, Peak Left Ventricular Pressure (Pmax), dP/dt Max (dP/dt+), dP/dt Min (dP/dt-), Contractility Index (CI), Tau (T_1/2_), Heart Rate (bpm), Systolic Ejection Period (SEP), Diastolic Filling Period (DFP), Contraction Time (CT) and Relaxation Time (RT). For Systemic Blood Pressure: Systolic Blood Pressure (mmHg), Diastolic Blood pressure (mmHg), Mean Arterial Blood Pressure (mmHg) and Pulse Pressure (mmHg).

### Tissue harvest and collection

While under anesthesia (isoflurane) following terminal cardiovascular assessments, blood collection was performed using the catheter placed during cardiovascular assessment. At PPD120, serum was collected. For serum collection, blood was collected in serum separator tubes and kept on wet ice or ice packs until serum isolation, within 60 minutes of collection using a refrigerated centrifuge at room temperature for at least 10 minutes at a relative centrifugal force of at least 1800g. Once the serum was isolated, it was saved in aliquots, and each sample was flash frozen in liquid nitrogen within 2 hours of collection and stored in a freezer set to maintain -85 to -60°C. Following blood collection, while under anesthesia (isoflurane), animals were humanely euthanized via exsanguination. As soon as feasible following euthanasia, the heart was removed, weighed, rinsed with Phosphate Buffered Saline (PBS) and sectioned. The heart was bisected along its longitudinal axis parallel to the interventricular septum (IVS), with the IVS retained with the right side of the heart. The apex of the heart was removed from the left side and flash frozen. The remainder of the left side was flash frozen. The samples were maintained on dry ice until stored in a freezer set to maintain -85 to -60°C. The right side of the heart was emersed in NBF for at least 24 hours. After 24 hours in NBF, the heart was removed, and a thin slice of the right ventricular free wall was transferred to glutaraldehyde and processed for µultrastructural analysis. The remainder of the right side of the heart was returned to NBF until fully fixed and processed for histopathology. For animals removed at 13 weeks after weaning, following collection of the heart samples, two samples (∼250 mg total) from the left liver lobe were collected, cut into smaller pieces (approximately 5 mm3), and flash frozen. The samples were maintained on dry ice until stored in a freezer set to maintain -85 to -60°C.

### Human induced pluripotent stem cell-derived cardiomyocyte culture and MEA

Cryopreserved hiPSC-CM (iCell Cardiomyocyte^2^) were obtained from FUJIFILM Cellular Dynamics (FCDI, Madison, WI). These cells were plated with iCell Cardiomyocytes Plating Medium (FCDI) and maintained in iCell Cardiomyocyte Maintenance Medium (FCDI) according to manufacturer’s protocols. For multielectrode array, cells were plated on Axion Biosystems CytoView MEA Plates (Axion BioSystems, USA) according to manufacturer’s protocols. Cells were treated with ABC/DTG/3TC at concentrations equivalent to 50% of each respective drug concentration in our in vivo model, equating to 8µg/mL / 9µg/mL / 2µg/mL, respectively. Cells were treated every 48 hours in iCell Maintenance Medium. After 7 days of treatment, cell functionality was analyzed via multielectrode array using Maestro Pro MEA platform (Axion Biosystems). Cardiac field potential recordings were performed using the Maestro Pro MEA platform (Axion BioSystems) at 37 °C and 5% CO2. Real-time data was collected simultaneously across all 768 electrodes (12.5 kHz sampling rate, 1-2000 Hz band-pass). Analysis was performed using AxIS Navigator software and the Cardiac Analysis Tool (Axion BioSystems). Local extracellular action potential (LEAP) recordings were performed using the Maestro Pro MEA platform (Axion BioSystems) at 37 °C and 5% CO2. Real-time data was collected simultaneously across all 768 electrodes (12.5 kHz sampling rate, 0.01-2000 Hz band-pass). LEAP induction was performed by enhancing cell coupling through electrical stimulus for 10 minutes using the AxIS Navigator software (Axion BioSystems). Analysis was performed using AxIS Navigator and the Cardiac Analysis Tool software (Axion BioSystems). After measurement, cells were harvested and snap frozen for further analysis.

### DNA isolation, Mitochondrial copy number and deletions

DNA was isolated from heart and liver tissue and hiPSC-CMs using DNeasy Blood and Tissue Kit (QIAGEN, Valencia, CA) following manufacturer protocol. DNA concentration was estimated in NanoDrop^TM^ 2000 and Qubit^TM^ 3 fluorometer. After isolation, DNA was diluted in DEPC water to a concentration of either 0.5ng/µL (hiPSC-CM), 0.25 ng/µL (liver) or 0.05 ng/µL (heart) for qPCR analysis of mitochondrial genes as surrogates for mitochondrial copy number and mitochondrial genome deletions. PCR amplification was performed using primers for *18S, ND1, ND4L, CYTB, COX2* and *ND3*. Sequences for these primers can be found in the supplement (Fig. S4). PCR amplification was performed using CFX384 Real-Time PCR System (Bio-Rad, Hercules, CA) and analyzed using primer efficiencies. 7S DNA level was quantified as previously described ^42^, adjusting the primers to the R. norvegicus mtDNA sequence. Briefly, ΔCt was calculated as the difference between the Ct corresponding to primers amplifying the 7S and mtDNA (7S-A and 7S-B1) and Ct corresponding to primers amplifying mtDNA only (7S-A and 7S-B2) (Fig. S4), followed by normalization to the average value of the PPD21 control samples. The formula 2^(−ΔΔCt) was used to calculate the fold enrichment.

### Oxygen consumption rate (OCR) in hiPSC-CM

OCR in intact cells was performed as previously described ^43^. Briefly, OCR was assessed using the Seahorse XFe96 Analyzer (Agilent Technologies). Seahorse XFe96 Cell Cultured Microplates (Agilent Technologies) were coated with 50µg/mL fibronectin and seeded with hiPSC-CM (10,000 cells/well) following manufacturer’s protocol (FIDC) and were incubated at 37 °C in a 5% CO_2_ humidified incubator. Cells were treated as previously described with ABC/DTG/3TC in the cell culture microplate every other day for 7 days, at which point they were washed twice by dilution with supplemented XF Assay medium and incubated in a O_2_-free BOD-type incubator at 37 °C for 1 h. MitoStress protocol was used to access mitochondrial function. At the end of the assay, the medium was removed, and cells were washed with PBS. PBS was removed and cells were lysed with 20µL of RIPA Lysis and Extraction Buffer supplemented with Halt™ Protease and Phosphatase Inhibitor Cocktail (Thermo Scientific^TM^). Protein content was estimated using Pierce™ BCA Protein Assay Kits (ThermoFisher™). OCR values were corrected by protein content of each given well. The average of 6–8 replicates was considered one independent sample. OCR and ECAR results were transformed in O_2_ pmol/min/mg protein and mpH/min/mg.

### OCR in frozen tissue

OCR was assessed in frozen placental lysate in a Seahorse XFe96 Analyzer (Agilent Technologies). Our group adapted a protocol that has been previously published detailing how to measure mitochondrial respiration in frozen biological samples^18^. Briefly, 20mg of tissue was homogenized in physiologic buffer (70mM sucrose, 220mM mannitol, 5mM KH2PO4, 1mM EGTA, 2mM HEPES) using an OMNI homogenizer with a soft tissue attachment (Omni International). Samples were spun down to enrich for a mitochondrial fraction and supernatant was collected. Protein content was estimated using Pierce BCA Protein Assay Kits (ThermoFisher) and used to standardize content across wells. Protein diluted in physiologic buffer was plated in Seahorse XFE96 Cell Culture Microplates. Before running, 115 µL of physiologic buffer + 11.8 µg/mL cytochrome C was added to every well. Final concentration of substrates and inhibitors was: NADH 1mM or Succinate 5mM + Rotenone 2µM (Port A), Antimycin A (Port B), TMPD 0.5mM + Ascorbate 1mM (Port C), and Sodium Azide 50mM (Port D) (all Sigma-Aldrich). OCR values were corrected by TOMM20 values as determined by the ProteinSimple Jess System (Bio-Techne). The average of 6-8 replicates from the original lysate was considered one independent sample (n=6). OCR results were transformed in O2 pmol/min/TOMM20.

### Transmission Electron Microscopy of Cardiac Tissue

Electron microscopy of cardiac tissue was performed as previously described^43^. Briefly, cardiac sections were fixed for 1 hour in fixative and washed twice in 0.1M cacodylate buffer prior post-fixation in osmium tetroxide 1%. Tissue was rinsed in distilled water and dehydrated in an ethanol series which transitioned to acetone. Samples were infiltrated with resin and polymerized, and blocks were cut, mounted on glass slides and stained in 1% toluidine blue in 1% sodium borate for quality check examination with a light microscope. After trimming, ultrathin sections were cut from blocks, placed in blocks and counterstained with 2% uranyl acetate and lead citrate. Using an FEI Co. Tecnai T12 transmission electron microscope operated at 80kV, slides were visualized and digitally captured. For quantification, at least 100 images for each study group were captured and EM subcellular analysis was made using Gatan Digital Micrograph software.

### RNA isolation, RNA-seq and Gene Expression Analysis

RNA was extracted from frozen cardiac or liver tissue for RNA-seq using the RNeasy Kit (QIAGEN). The total RNA concentration was determined using the Qubit Flex Fluorometer (Thermo Fisher Scientific) and the RNA integrity was analyzed using an Agilent’s TapeStation 4200. RNASeq libraries were created using Illumina’s TruSeq Starnded mRNA kit and verified using an Agilent TapeStation 4200. The RNAseq libraries were sequenced on Illumina’s NovaSeq platform at a read length of 75bp and with a single end format. Single-end 76-cycle reads were aligned to the rat rn7 reference genome with STAR version 2.6. An average of 52 and 44 million uniquely mapped reads were obtained per heart and liver sample, respectively. Raw read counts were obtained using Rsubread version 2.14. Normalized counts and differentially expressed genes were obtained using DESeq2 version 1.42. Heatmaps and 3D PCA plots were created using R packages pheatmap version 1.0 and rgl version 1.3, respectively. For age-related comparisons, PPD120 heart samples were compared to PPD21 heart samples in the control, low dose and high dose groups. For treatment comparisons, livers at PPD120 were compared using vehicle control as the baseline vs low dose- or high dose-treated animals. Genes with a fold change ≥ 1.5 and FDR < 0.05 were considered statistically significant.

### Metabolomics

Metabolomic profiling was completed on hearts from dams at PPD21 and PPD120 and stored at −80 °C. Equal volumes of all samples were extracted and run across the Metabolon Inc. (Durham, NC) Precision Metabolomics discovery platform. Samples were split into equal parts for analysis on four LC/MS/MS platforms. All methods utilized a Waters ACQUITY ultra-performance liquid chromatography (UPLC) and a Thermo Scientific Q-Exactive high resolution/accurate mass spectrometer interfaced with a heated electrospray ionization (HESI-II) source and Orbitrap mass analyzer operated at 35,000 mass resolution ^44^. Proprietary software was used to match to an in-house library of standards for metabolite identification and for metabolite quantitation by peak area integration, where a metabolite’s abundance was defined as its peak area. Peaks were quantified using area-under-the-curve. Metabolon data analysts use proprietary visualization and interpretation software to confirm the consistency of peak identification and integration among the various samples.

### Clinical Chemistry

The following clinical chemistry parameters were analyzed using a Roche Cobas c311 chemistry analyzer (Indianapolis, IN): total cholesterol, low-density lipoprotein (LDL) cholesterol, high-density lipoprotein (HDL) cholesterol and triglycerides. The following clinical chemistry parameters were analyzed using a Beckman Coulter AU480 chemistry analyzer (Brea, CA): alkaline phosphatase (ALP), alanine aminotransferase (ALT), aspartate aminotransferase (AST), total bilirubin, direct bilirubin and total bile acids.

### Statistical Analysis

All data was analyzed using GraphPad Prism (La Jolla, CA). Statistical outliers were identified using the ROUT method (Q = 1%). When comparing two groups, an unpaired Student’s t test was used. Statistical comparisons between multiple groups were determined using an ordinary one-way or two-way ANOVA with multiple comparisons test as appropriate. All tests were two-sided. Significance was set for all experiments at p < 0.05.

## Acknowledgements

We would like to thank the staff from the Epigenomics Core Facility at NIEHS for their technical support. We thank Drs. Vesna Chappell and G. Jean Harry (NIEHS) for fruitful discussions. We also thank Drs. Andreas Beyer (Medical College of Wisconsin) and June Dunnick (NIEHS) for critical review of the manuscript.

## Author Contributions

Conceptualization: JHS

Methodology: GR, VS, JHS, HC, MPM, CAN, NT, SC

Investigation: NT, MPM, CAN, SC, JH, MJ, GLC

Formal Analysis: NT, MJ, JH, MPM, SC, CAN, GLC

Visualization: NT, JHS

Supervision: MC, SW, GR, JHS

Writing—original draft: NT, JHS Writing—review & editing: NT, JHS

## Data and materials availability

The RNA-seq and metabolomics data generated in this study will be deposited in the GEO database prior to publication. Source data will be provided with this paper prior to publication.

