## Supplementary material for "Antiretroviral therapy in the peripartum period impairs post pregnancy cardiac reverse remodeling with early signs of heart failure with preserved ejection fraction": suppl legends

**Supplemental Figure Legends**

**Supplemental Table 1. RNA-seq from heart tissue from exposed and non-exposed Sprague-Dawley female rats.**

**Supplemental Table 2. Metabolomics analysis of heart tissue derived from vehicle- or cART-exposed Sprague-Dawley female rats.**

**Supplemental Table 3. RNA-seq from liver tissue from exposed and non-exposed Sprague-Dawley rats.**

**Supplemental Fig. 1. Fertility and fecundity parameters and gross organ pathology from cohorts of Sprague-Dawley female rats exposed to two different doses of ABC/DTG/3TC for 5.5 months.**

**Supplemental Fig. 2. Cardiac parameters evaluated in dams administered ABC/DTG/3TC.** (A) Cardiac parameters shown as a function of time where repeated measures on the same animals evaluated heart rate (HR), cardiac output (CO), stroke volume (SV), ejection fraction (EF), and fractional shortening (FS) (n=12 rats/group). Analysis performed via two-way ANOVA with multiple comparisons, p-values shown are corresponding group vs. vehicle control. (B) Dobutamine challenge results at both postpartum day 21 (PPD21) and postpartum day 120 (PPD120) (n= 12 rats/group). Increasing doses of dobutamine were infused per minute, which are depicted in the figure. Analysis performed via two-way ANOVA with multiple comparisons, p-values shown are corresponding group vs. vehicle control. (C) Blood pressure measurements at PPD21 and PPD120 (n=12 rats/group). (D) QRS interval as measured by ECG (n=12 rats/group).

**Supplemental Figure 3. Additional mitochondrial measurements for rats and iPSC cardiomyocytes administered ABC/DTG/3TC.** (A) Mitochondrial DNA genes amplified at PPD21 and PPD120 via qPCR, including *ND1, COX2, CYTB, ND4L* and *18S* (n=12 rats/group). (B) Mitochondrial respiration in iPSC cardiomyocytes exposed to either media, DMSO 0.02% or ABC/DTG/3TC for 7 days (n=6/group) using SeaHorse Flux Analyzer. (C) SDHA and TOMM20 protein levels in the hearts of rats exposed to low and high dose ABC/DTG/3TC measured by SimpleWestern Jess at PPD21 and PPD120 (n=12 hearts/group). (D) Electron microscopy in heart tissue of rats exposed to low and high dose ABC/DTG/3TC; data are representative of n=2/dose/timepoint with 2 independent blocks analyzed per animal. About 100 images were taken/block.

**Supplemental Figure 4. qPCR primer sequences used for mitochondrial DNA copy number and 7S DNA.**
