## Supplementary material for "Antiretroviral therapy in the peripartum period impairs post pregnancy cardiac reverse remodeling with early signs of heart failure with preserved ejection fraction": suppl data

### Supplementary Data 1

## A

| Sex: Female | | 0 mg/kg | 150/12.5/ 75 mg/kg | 300/25/ 150 mg/kg |
|---|---|---|---|---|
| Group Size | | 34 | 33 | 33 |
| Pregnant? | N+ve | 25 | 27 | 27 |
| Fertility | | 0.735 | 0.818 | 0.818 |
| Littered | N+ve | 24 | 26 | 26 |
| Viable | N+ve | 24 | 26 | 26 |
| Fecundity | | 0.960 | 0.963 | 0.963 |
| Gestation Length NTP | Mean | 22.3 | 22.2 | 22.4 |
| | SEM | 0.1 | 0.1 | 0.1 |
| | N | 24 | 26 | 26 |
| | %Diff | . | -0.3 | 0.6 |

## B

| Removal Reason(s): ALL<br>Summary: Average Grade (Mean) | Female<br>0 mg/kg | <br>150/12/75 mg/kg | <br>300/25/15 mg/kg |
|---|---|---|---|
| Number of Animals: | 12 | 12 | 12 |
| Number of Completed Animals: | 12 | 12 | 12 |
| **liver** | | | |
| Examined | 12 | 12 | 12 |
| No Visible Lesions | 12 | 12 | 11 |
| periportal; infiltrate, mononuclear cell | 0 | 0 | 1 |
| .... minimal | 0 | 0 | 1 |
| .... Average Grade | 0.0 | 0.0 | 1.0 |
| **lung** | | | |
| Examined | 12 | 12 | 12 |
| No Visible Lesions | 0 | 2 | 3 |
| subpleura; interstitium; inflammation; chronic | 8 | 8 | 7 |
| .... minimal | 6 | 7 | 6 |
| .... mild | 2 | 1 | 1 |
| .... Average Grade | 1.3 | 1.1 | 1.1 |
| alveolus; infiltration cellular; histiocyte | 10 | 3 | 4 |
| .... minimal | 7 | 3 | 3 |
| .... mild | 3 | 0 | 1 |
| .... Average Grade | 1.3 | 1.0 | 1.3 |
| blood vessel; infiltration cellular; eosinophil, perivascular | 2 | 0 | 0 |
| .... minimal | 2 | 0 | 0 |
| .... Average Grade | 1.0 | 0.0 | 0.0 |
| metaplasia; osseous | 1 | 0 | 0 |

B

### Supplementary Data 1

| Removal Reason(s): ALL<br>Summary: Average Grade (Mean) | Female<br>0 mg/kg | Female<br>150/12/75 mg/kg | Female<br>300/25/15 mg/kg |
|---|---|---|---|
| Number of Animals: | 12 | 12 | 12 |
| Number of Completed Animals: | 12 | 12 | 12 |
| **lung (Continued...)** | | | |
| blood vessel; mineral | 1 | 1 | 0 |
| .... minimal | 1 | 1 | 0 |
| .... Average Grade | 1.0 | 1.0 | 0.0 |
| **diaphragm** | | | |
| Examined | 12 | 12 | 12 |
| No Visible Lesions | 12 | 12 | 12 |
| **mesenteric roll** | | | |
| Examined | 12 | 12 | 12 |
| artery 1; no visible lesions | 9 | 8 | 8 |
| artery 2; no visible lesions | 10 | 9 | 8 |
| artery 3; no visible lesions | 11 | 10 | 9 |
| artery 4; no visible lesions | 12 | 10 | 9 |
| artery 5; no visible lesions | 12 | 10 | 9 |
| artery 1; mineral | 3 | 4 | 4 |
| .... minimal | 3 | 4 | 4 |
| .... Average Grade | 1.0 | 1.0 | 1.0 |
| artery 2; mineral | 2 | 3 | 4 |
| .... minimal | 2 | 3 | 4 |
| .... Average Grade | 1.0 | 1.0 | 1.0 |
| artery 3; mineral | 1 | 2 | 3 |

| Removal Reason(s): ALL<br>Summary: Average Grade (Mean) | Female<br>0 mg/kg | Female<br>150/12/75 mg/kg | Female<br>300/25/15 mg/kg |
|---|---|---|---|
| Number of Animals: | 12 | 12 | 12 |
| Number of Completed Animals: | 12 | 12 | 12 |
| **mesenteric roll (Continued...)** | | | |
| .... minimal | 1 | 2 | 3 |
| .... Average Grade | 1.0 | 1.0 | 1.0 |
| artery 4; mineral | 0 | 2 | 3 |
| .... minimal | 0 | 2 | 3 |
| .... Average Grade | 0.0 | 1.0 | 1.0 |
| artery 5; mineral | 0 | 2 | 3 |
| .... minimal | 0 | 2 | 3 |
| .... Average Grade | 0.0 | 1.0 | 1.0 |

Supplementary Data 2

A

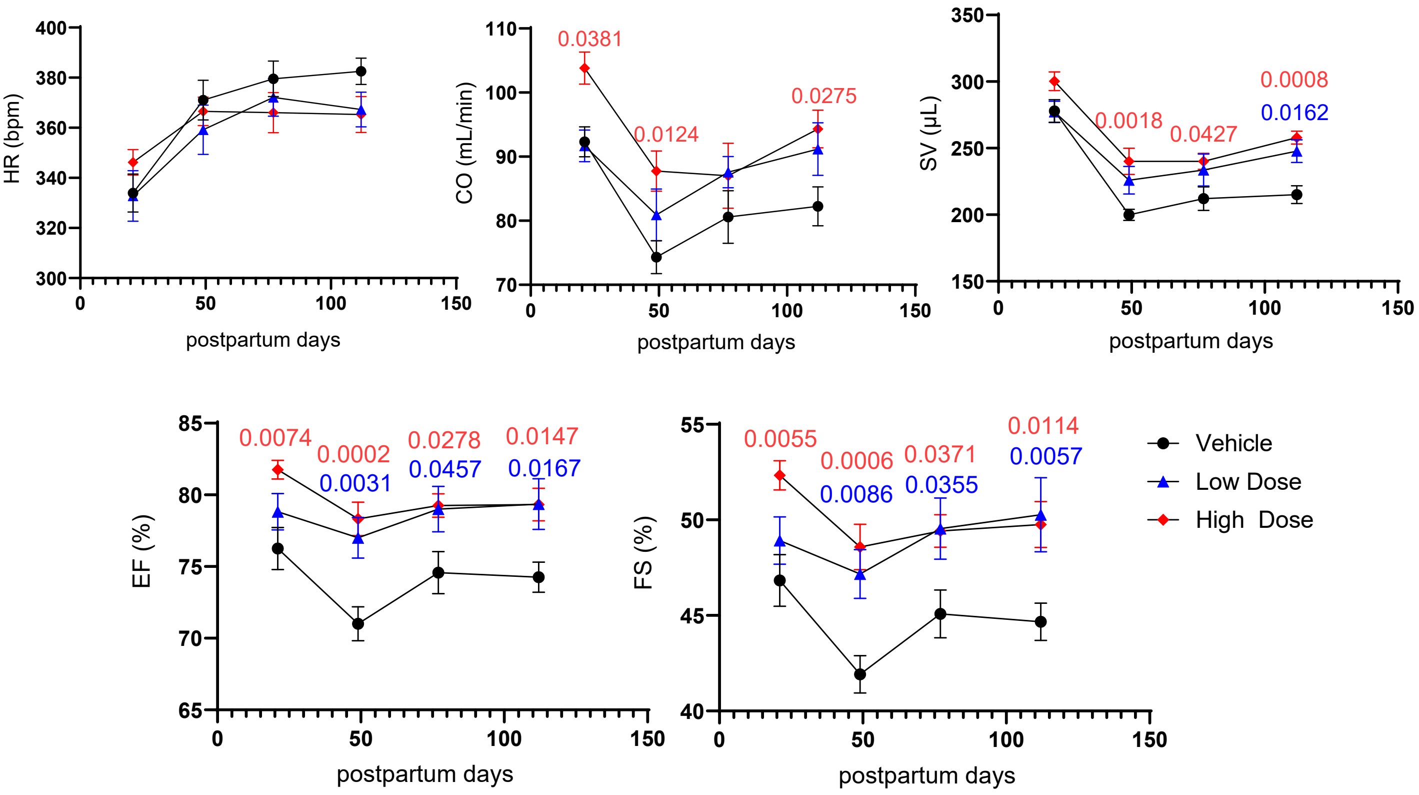


B

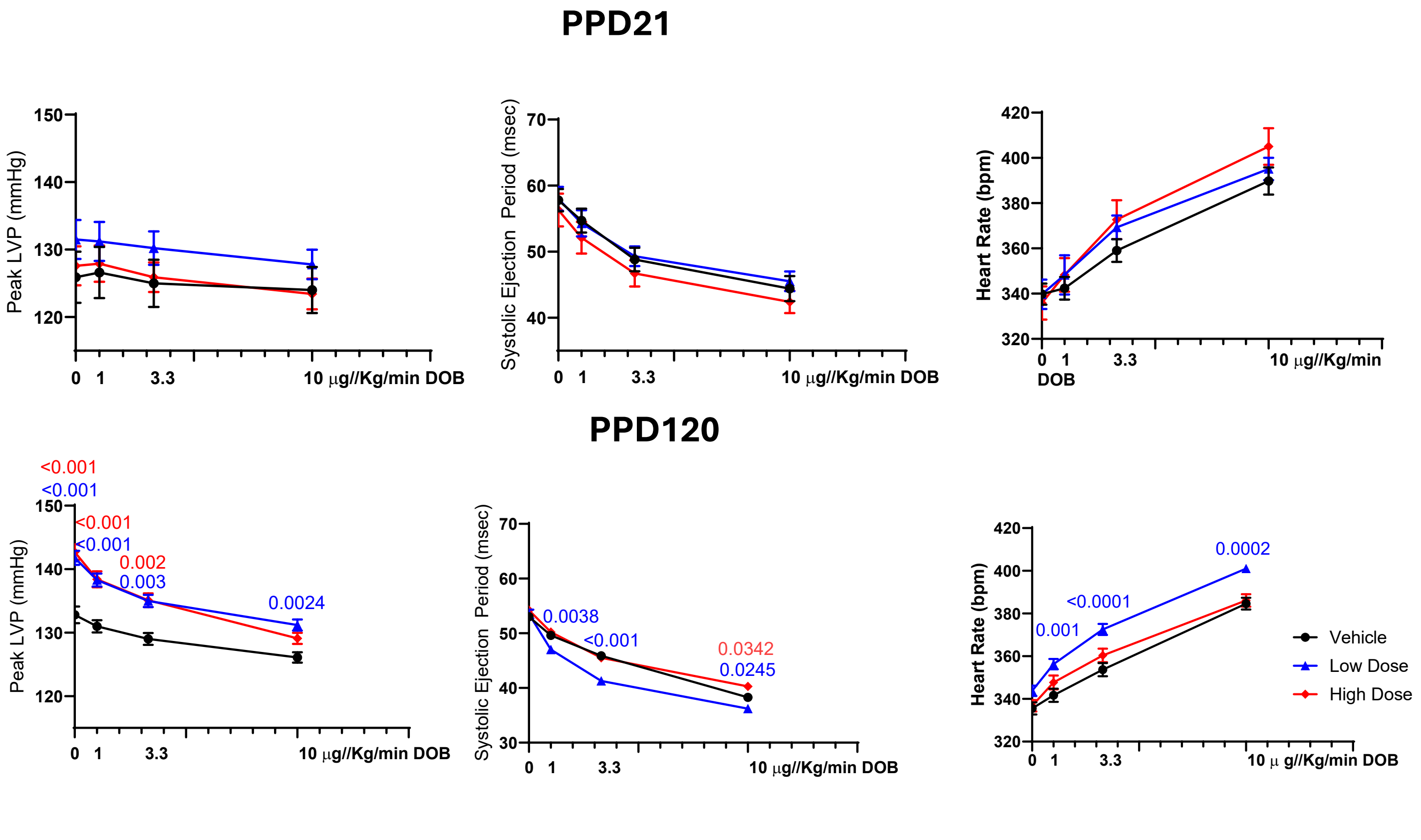

Supplementary Data 2

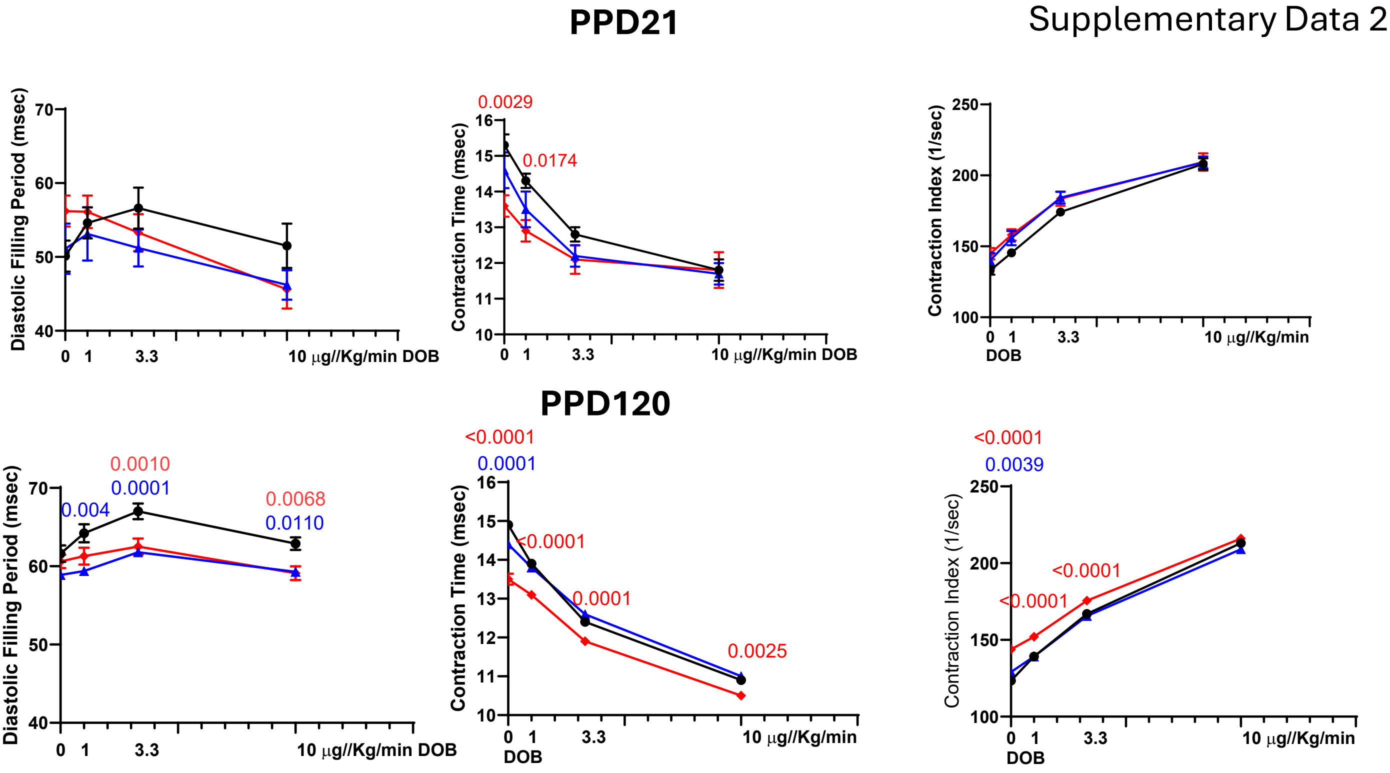


C

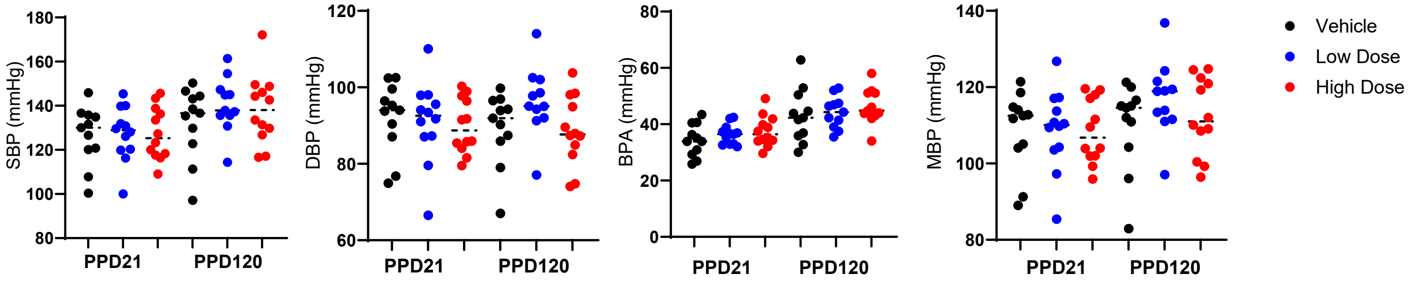


D

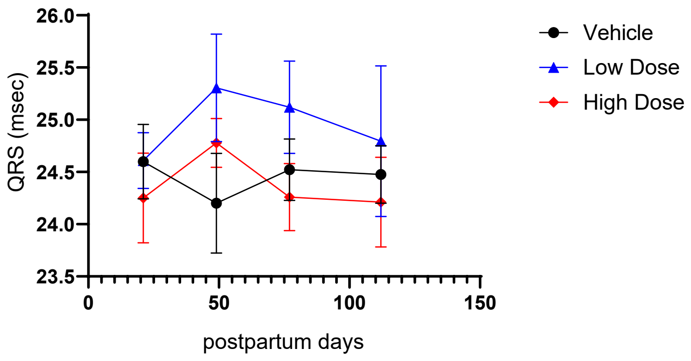

Supplementary Data 3

A

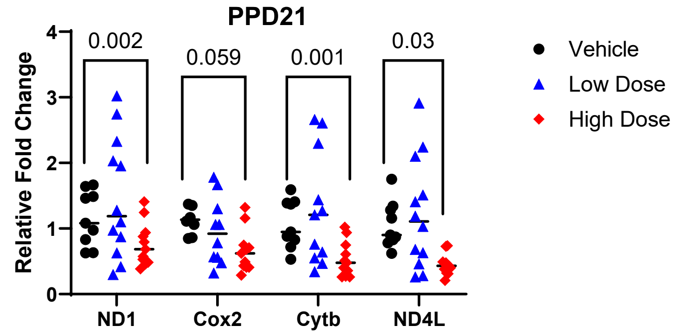


B

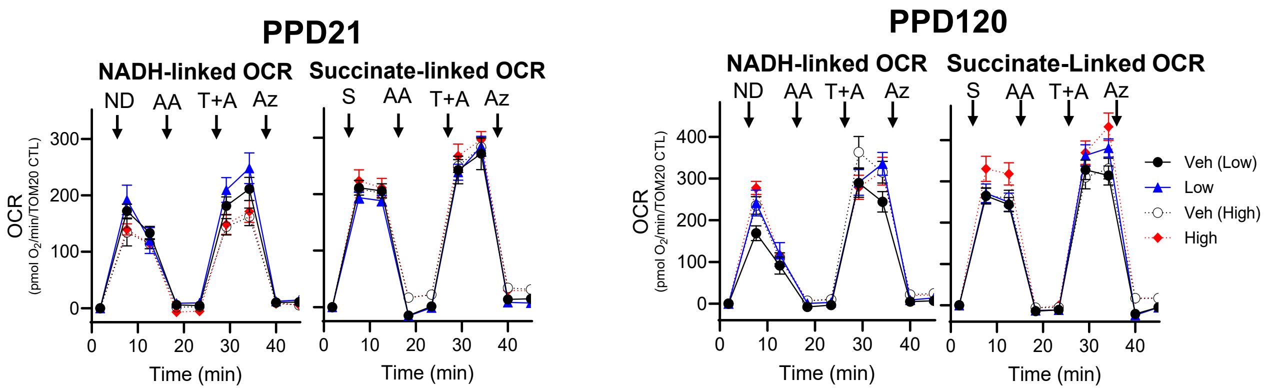


C

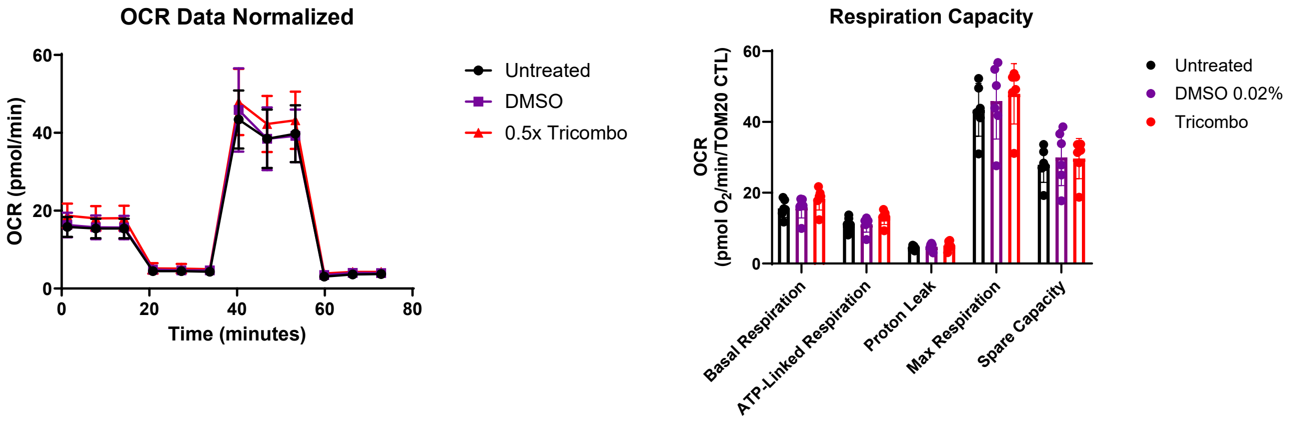


D

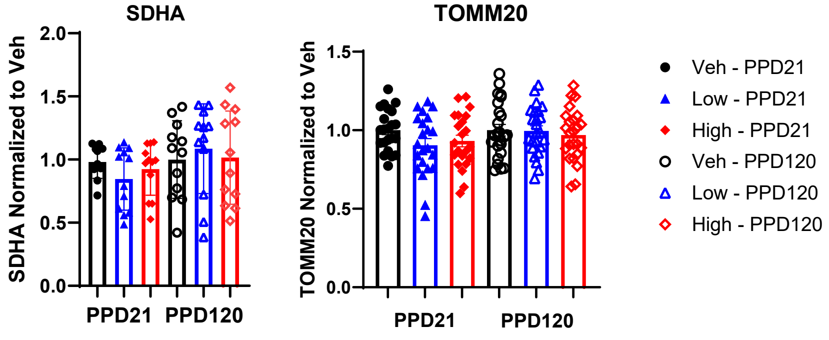

Supplementary Data 3

D

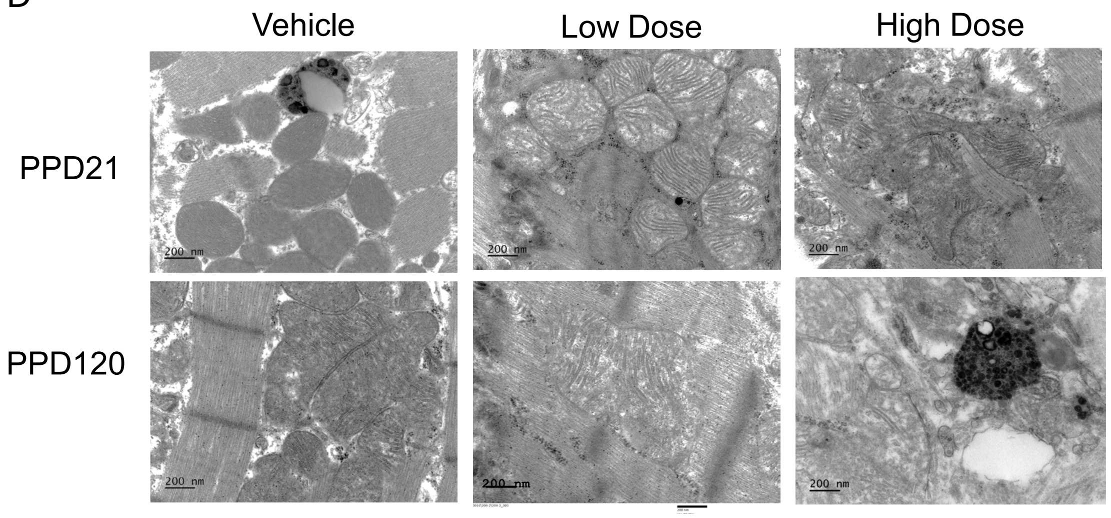

| Gene Name | Organism | F1 Sequence | R1 Sequence |
|---|---|---|---|
| 18S | Rat | TCAACTTTCGATGGTAGTCGCC | TTGGATGTGGTAGCCGTTTCT |
| ND1 | Rat | TTCCTAGGCCCCTTATATCACA | ATCGAAAACGGGGGTAGGATG |
| ND4L | Rat | TGCGAAGCAGCAGTAGGTTTA | TTTGTACGTAGTCTGTTCCGT |
| CYTB | Rat | CCTCCCATTCATTATCGCCGCC CTTGC | GTCTGGGTCTCCTAGTAGGTCTG GGAA |
| COX2 | Rat | ATCCGAAGACGTCCTCGACT | ACTGTAGCTTGGTTTAGGCGG |
| ND3 | Rat | CCCATATGAATGTGCCTTCGAC | TTG AAA AAG GAA GGC GTG CAG |
| 7S-A | Rat | GAGGATGGTAGAAATAGAGACC | |
| 7S-B1 | Rat | CCCCAAGCATATAAGCATGTAA TA | |
| 7S-B2 | Rat | ACCATCAACACCCAAAGCTG | |
| 18S | Human | CGGACAGGATTGACAGATT | CCAGAGTCTCGTCGTTATCG |
| ND1 | Human | CGATTCCGCTACGACCAACT | AGGTTTGAGGGGGAATGCTG |
| ND4L | Human | ATCGCTCACACCTCATATCCT | GGAGTGGGTGTTGAGGGTTAT |
